# nP-collabs: Investigating counterion mediated bridges in the multiply phosphorylated tau-R2 repeat

**DOI:** 10.1101/2024.04.18.590060

**Authors:** Jules Marien, Chantal Prévost, Sophie Sacquin-Mora

## Abstract

Tau is an instrinsically disordered (IDP), microtubule-associated protein (MAP) that plays a key part in microtubule assembly and organization. The function of tau can be regulated via multiple phosphorylation sites. These post-translational modifications are known to decrease the binding affinity of tau for microtubules, and abnormal tau phosphorylation patterns are involved in Alzheimer’s disease. Using all-atom molecular dynamics (MD) simulations, we compared the conformational landscapes explored by the tau R2 repeat domain (which comprises a strong tubulin binding site) in its native state and with multiple phosphorylations on the S285, S289 and S293 residues, with four different standard force field (FF)/water model combinations. We find that the different parameters used for the phosphate groups (which can be more or less flexible) in these FFs, and the specific interactions between bulk cations and water lead to the formation of a specific type of counterion bridge, termed *nP-collab* (for nPhosphate collaboration, with *n* being an integer), where counterions form stable structures binding with two or three phosphate groups simultaneously. The resulting effect of nP-collabs on the tau-R2 conformational space differs when using sodium or potassium cations, and is likely to impact the peptide overall dynamics, and how this MAP interacts with tubulins. We also investigated the effect of phosphoresidues spacing and ionic concentration by modeling polyalanine peptides containing two phosphoserines located one to six residues apart. Three new metrics specifically tailored for IDPs (Proteic Menger Curvature, Local Curvature and Local Flexibility) were introduced, which allow us to fully characterize the impact of nP-collabs on the dynamics of disordered peptides at the residue level.

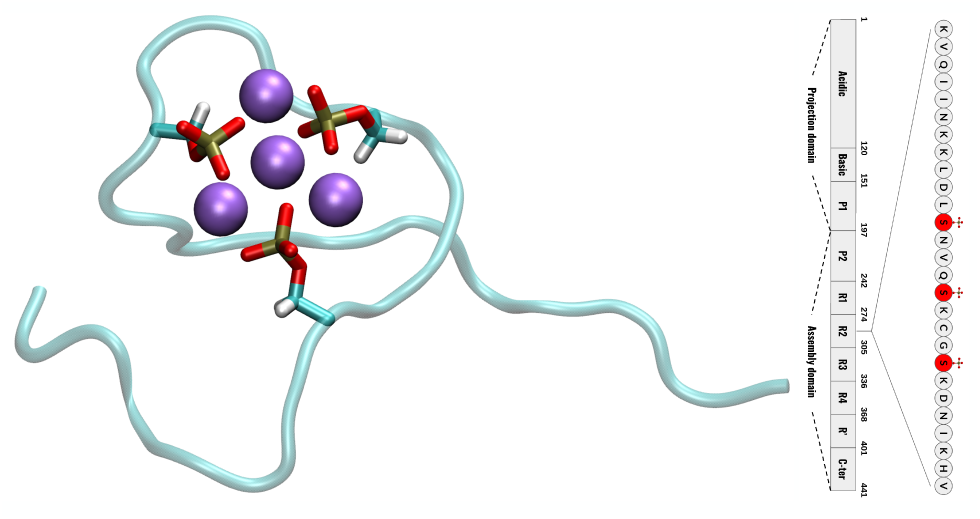

## Introduction

Tau is a Microtubule Associated Protein (MAP) abundant in neuronal tissues,^1, 2^ which plays a key part in the regulation of microtubule (MT) assembly and spatial organization, and also controls the motility of motor proteins along MTs.^3–7^ The function of this member of the intrinsically disordered protein (IDP) group is regulated by post-translational modifications (PTM) such as phosphorylation, acetylation, glycosylation or ubiquitination.^8–10^ While MTs regulation relies on reversible tau phosphorylation in a healthy brain, tau hyper- phosphorylations have been shown to make tau loose its affinity for MTs, promote aggregation and be involved in several pathologies, including neurodegenerative diseases (tauopathies) such as Alzheimer’s disease (AD).^11^ For example, in AD brains, tau can be phosphorylated on roughly 80 positions (mostly serine and threonine residues),^12, 13^ and these PTMs are likely to disrupt the interaction of tau with MTs and the stabilization of the MT filament.^14–16^ More generally, phosphorylations (mostly located on serine and threonine residues) are the most common PTM in mammalians^17^ and are involved in numerous cellular processes and human diseases (like neurodegenerative disorders or cancers).^18–20^ IDPs are a common target for phosphorylations, which impact their charge, folding propensity, dynamics and binding properties, with important consequences for their biological function.^21–23^

From the experimental point of view, one can resort to several techniques, such as nuclear magnetic resonance (NMR), circular dichroism (CD), small-angle X-ray scattering (SAXS) or Förster resonance energy transfer (FRET) to obtain information regarding the impact of phosphorylations on an IDP’s conformational ensemble and mechanical properties.^24–29^ These approaches are also well complemented by atomistic simulations, which bring further insight on the conformational space covered by the protein, and the last decade has seen the development of numerous force-fields that were specifically parametrized for the modeling of disordered proteins.^30–35^

Following an earlier study using molecular dynamics simulation to investigate the interaction pattern in the *fuzzy complex*^36^ formed by the tau-R2 fragment (which comprises a strong interacting region centered around Ser289^37^) and the tubulin surface,^38^ we phosphorylated the Ser285, Ser289 and Ser293 residues in the isolated tau-R2 fragment, which have been identified as phosphorylation sites in tauopathies.^11, 14–16^ Ser289, in particular, is one the 40 phosphorylations sites found in AD brains, but not in healthy ones.^39^ Initial analyses of the conformational space explored by the multiply phosphorylated tau-R2 fragment highlighted stable structures with one or several Na^+^ counterions bridging two or three phosphate groups, which we named *nP-collabs* (for nPhosphate-collaborations, with *n* being an integer). In order to assess the quality of our simulation parameters and the reproducibility of this phenomenon, we decided to perform similar calculations on the isolated R2 fragment with four different force-field/water model combinations commonly used to model phosphorylations in proteins. The comparison of the resulting trajectories shows that nP-collabs are not specific to the Charmm36m force-field with which they were initially observed, but their formation depends on the combination of the parameters used for the phosphate groups, the bulk cations and the water molecules. Based on additional simulations, we also investigated the impact of the counterion nature (Na^+^ or K^+^) and concentration, and the position of the phosphorylation sites along the protein sequence on nP-collab formation using three new metrics : Protein Menger Curvature, Local Curvature and Local Flexibility. These metrics provide a promising framework for rigidity analysis in the larger context of IDPs. Finally, we discuss how nP- collabs are likely to impact the interaction of phosphorylated IDPs with their protein partners, and more specifically the tau/MT interface.

## Materials and Methods

### Simulations of the tau-R2 repeat in solution

We used the structure of the tau-R2 repeat (a 27 residue fragment ranging from Lys274 to Val300) extracted from the tau-tubulin assembly modeled in our previous work^38^ as a starting point. Phosphorylations were added using the CHARMM-GUI^40^ (https://www.charmm-gui.org/) input generator. Classical, all-atom, MD simulations were performed using GROMACS v2021.3 in a box of 10^3^ nm^3^ with a physiological solution (0.15 mol.L^-1^) of NaCl or KCl. Table 1 gives the main simulation details, such as the numbers of ions, water molecules and the phosphorylation states. The amino-acid protonation state was set to pH 7 (and histidine residues were unprotonated), leading to a +4 net charge in the unphosphorylated peptide. Upon phosphorylation, each phosphate group was chosen to be unprotonated and adds a charge of -2 to the peptide, as the most common form of the phosphate group at pH 7 is dianionic.^41^

**Table 1:** Summary of the tau-R2 MD simulations performed in the study. For each setup (column), we ran three replicas of 520 ns.

| FF/water combination | Cation | Phosphorylated serines | Protein atoms | Water molecules | Cations | Anion |
| --- | --- | --- | --- | --- | --- | --- |
| **C36** | Na<sup>+</sup> | no | 446 | 32431 | 89 | 93 |
| **C36** | Na<sup>+</sup> | S285, S289, S293 | 455 | 32425 | 91 | 89 |
| **C36** | Na<sup>+</sup> | S285 | 449 | 32430 | 89 | 91 |
| **C36** | Na<sup>+</sup> | S289 | 449 | 32431 | 89 | 91 |
| **C36** | Na<sup>+</sup> | S293 | 449 | 32428 | 89 | 91 |
| **C36** | Na<sup>+</sup> | S285, S289 | 452 | 32427 | 90 | 90 |
| **C36** | Na<sup>+</sup> | S285, S293 | 452 | 32428 | 90 | 90 |
| **C36** | Na<sup>+</sup> | S289, S293 | 452 | 32427 | 90 | 90 |
| **C36** | K<sup>+</sup> | no | 446 | 32431 | 89 | 93 |
| **C36** | K<sup>+</sup> | S285, S289, S293 | 455 | 32425 | 91 | 89 |
| **C36** | K<sup>+</sup> | S285 | 449 | 32430 | 89 | 91 |
| **C36** | K<sup>+</sup> | S289 | 449 | 32431 | 89 | 91 |
| **C36** | K<sup>+</sup> | S293 | 449 | 32428 | 89 | 91 |
| **C36** | K<sup>+</sup> | S285, S289 | 452 | 32427 | 90 | 90 |
| **C36** | K<sup>+</sup> | S285, S293 | 452 | 32428 | 90 | 90 |
| **C36** | K<sup>+</sup> | S289, S293 | 452 | 32427 | 90 | 90 |
| **A99** | Na<sup>+</sup> | no | 446 | 32489 | 89 | 93 |
| **A99** | Na<sup>+</sup> | S285, S289, S293 | 455 | 32477 | 91 | 89 |
| **A99** | K<sup>+</sup> | no | 446 | 32489 | 89 | 93 |
| **A99** | K<sup>+</sup> | S285, S289, S293 | 455 | 32477 | 91 | 89 |
| **A19** | Na<sup>+</sup> | no | 446 | 32423 | 89 | 93 |
| **A19** | Na<sup>+</sup> | S285, S289, S293 | 455 | 32421 | 91 | 89 |
| **A19** | K<sup>+</sup> | no | 446 | 32423 | 89 | 93 |
| **A19** | K<sup>+</sup> | S285, S289, S293 | 455 | 32421 | 91 | 89 |
| **A18** | Na<sup>+</sup> | no | 446 | 32458 | 89 | 93 |
| **A18** | Na<sup>+</sup> | S285, S289, S293 | 455 | 32446 | 91 | 89 |
| **A18** | K<sup>+</sup> | no | 446 | 32458 | 89 | 93 |
| **A18** | K<sup>+</sup> | S285, S289, S293 | 455 | 32446 | 91 | 89 |

We used four different protein force-field/water model combinations: i) CHARMM36m^42^ and the associated modified TIP3P model^42^, ii) AmberFF19SB^43^ with OPC^44^, iii) AmberFF99SB- ILDN^45^ with TIP4PD^46^, iv) AmberFB18^47^ with TIP3P-FB.^48^ For simplicity, these force-field/ water model combinations will be termed C36, A19, A99 and A18 systems respectively in the *results* and *discussions* parts of the article. The ionic parameters were taken from Beglov and Roux^49^ for Charmm36m and for AmberFF99SB-ILDN (standard parameters of Charmm22), from Sengupta et al.^50^ for AmberFF19SB (optimized for the OPC water model), and from the SPC/E set of Joung and Cheatham^51^ for AmberFB18 (as instructed in the publication of the force field development^47^).

For AmberFF99SB-ILDN, the phosphorylation force-field phosaa10^52^ was chosen with improvements by Steinbrecher et al.^53^. Its evolution, called phosaa19SB, was used for AmberFF19SB. AmberFB18 is actually the name of the phosphorylation force-field extending AmberFB15.^54^ CHARMM36m already contains phosphorylation parameters. Simulation parameters were chosen so as to correspond as closely as possible to the situation in the reference papers (except for Amber99SB-ILDN, where we followed the steps described by Rieloff and Skepö in ref. 29). A graphical summary of the force-field/water model/ionic parameters/phosphorylation terms used in this work is available in Figure SI-1. Additional information on how to obtain a proper GROMACS topology, the simulation start files, and a discussion of the chosen parameters are available in Supplementary Materials.

A leap-frog integrator with a timestep of 2fs was used to generate the trajectories. Frames were printed every 10ps (see discussion on the mobility of ions in the Supplementary materials for justification). Hydrogens were constrained using the LINCS algorithm.^55^ Electrostatic terms were calculated with PME with a Fourier spacing of 0.12 nm and an interpolation order of 4 for production. The thermostat was set to T = 300 K thanks to the v-rescale algorithm with a time constant of 0.1 ps for both protein and non-protein groups. Pressure was set to 1 bar with a Parrinello-Rahman barostat with isothermal compressibility 4.5x10^-5^ bar^-1^ and a time constant of 2 ps in the NPT ensemble.

CHARMM36m systems were first equilibrated for 2 ns in the NVT ensemble with positional harmonic restraints of 1000 kJ.mol^-1^.nm^-1^ applied to protein heavy atoms. Fourier spacing in that case was set to 0.16 nm.

For Amber systems, a 200 ps phase of equilibration was conducted in the NPT ensemble with the Berendsen thermostat, so as to bring the fluid density up to ∼1.00 kg.L^-1^ (more details are given in the Discussion on the simulation parameters in the Supplementary Information file). This step was followed by 2 ns in the NVT ensemble.

Restraints were then lifted for the production phase, for each setup listed in Table 1, we ran three replicas, with a production run of 520 ns each in the NPT ensemble. In total, an aggregate of 1.56 µs was obtained for every type of simulation. The complete input files are provided in a Zenodo repository (https://zenodo.org/records/12634115), and the convergence of the simulations (inter and intra-replica) is extensively discussed in the Supplementary Information file.

### Trajectory analysis

For the trajectory analysis and figures production, we used custom python scripts and the following packages or tools: MDAnalysis,^56^ MDTraj,^57^ Numpy,^58^ Matplotlib,^59^ Pandas,^60^ and VMD.^61^

To characterize the mechanical changes induced by nP-collabs in the peptide, we define three new metrics called Proteic Menger Curvature (PMC), Local Curvature (LC) and Local Flexibility (LF). The PMC is based on the Menger curvature^62^ applied to the proteic backbone. Let us imagine a triangle of summits A, B and C of area *A* and its circumcircle *C*.

The inverse of the radius of *C* is the Menger Curvature of the triangle. If we make B the α- carbon of residue *n*, A the α-carbon of residue *n-2* and C the α-carbon of residue *n+2*, we obtain a value of curvature that we call PMC*_n_* for residue *n*. We define the average of PMC*_n_* over the whole trajectory as the Local Curvature LC*_n_*, and the standard deviation as the Local Flexibility LF*_n_*. A schematic representation for the PMC, LC and FC is available in Figure SI-2. The python scripts for computing these metrics is available online (https://github.com/Jules-Marien/Articles/tree/main/_2024_Marien_nPcollabs/Demo_and_scripts_Proteic_Menger_Curvature). As these three metrics are based on internal coordinates, one should note that, unlike RMSD and RMSF, they do not require the alignement of the explored conformations along the trajectory.

## Results

### Conformational ensembles of the unphosphorylated tau-R2

We first investigated the conformational variability of unphosphorylated tau for the four considered force-field/water model combinations. The resulting distributions for the radius of gyration are shown in Figure 1a and 1c (left panels). The A18 simulations lead to slightly more compact structures of R2 compared to the other FF/water combinations (both with Na^+^ and K^+^ counterions), which can be related to the fact that unlike C36, A19 and A99, the A18 FF was not specifically developed for the modeling of IDPs. This is highlighted by higher LC values for the A18 system in the middle of the sequence compared to the rest of the force- fields (see the central panel for Figures 1a and 1c). In all cases, the fragment remains disordered during the simulations, as can be seen from the secondary structure propensities (shown in Figure SI-3 and SI-4), with only transient occurrences of α-helices and β-sheets.

**Figure 1:**
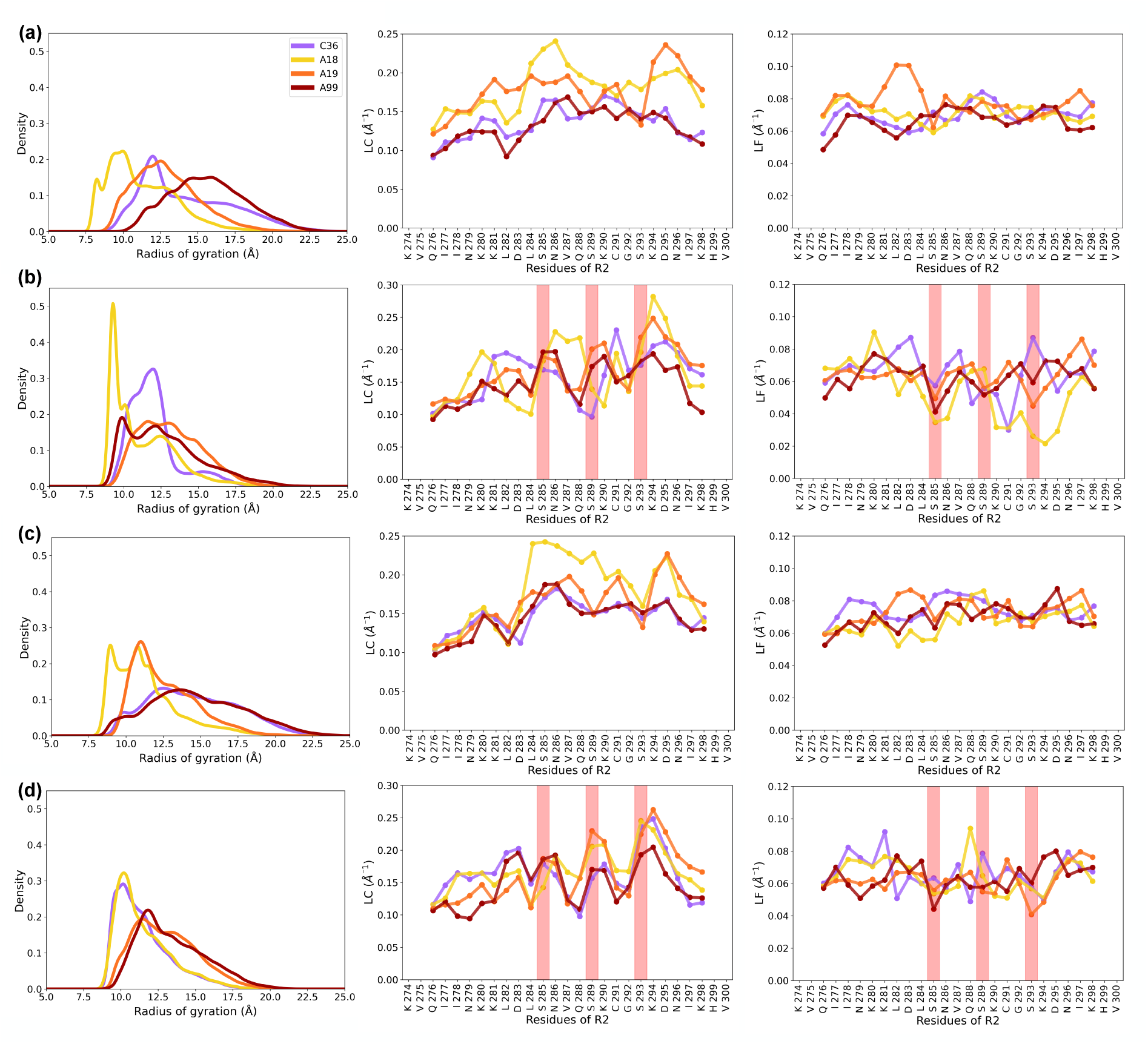
Comparison of the four considered FF/water model combinations. Left panels, density distribution of the radius of gyration of the tau-R2 fragment; central panels, local curvature (LC) along the sequence; right panels, local flexibility (LF) along the sequence. Simulations with Na^+^ counterions **(a)** Unphosphorylated tau **(b)** Triply phosphorylated tau. Simulations with K^+^ counterions **(c)** Unphosphorylated tau **(d)** Triply phosphorylated tau. The red columns in lines (b) and (d) highlight the positions of the phosphorylated serines along the sequence. LC and LF are obtained after concatenating the trajectories of the three replicas.

### Multiple phosphorylations of tau-R2 and nP-collab definition

The phosphorylation of serines S285, S289 and S293 in tau-R2 leads to a clear shift of the Rg distributions toward more compact structures for both counterion types (see the left panel in Figure 1b and 1d). This compaction of the peptide can be related to the formation of salt- bridges between the lysine residues of tau-R2 and the phosphate groups (see the averaged contact maps in Figure SI-5), a phenomenon commonly observed upon phosphorylation of positively charged IDPs.^21^ But during the MD trajectories, we also observed the formation of bent structures, where two or three phosphorylated serines are bridged by one or several positively charged counterion(s) (see the representative snapshot in Figure 2), in a similar fashion to the counterion bridges observed by Tolmachev et al. in polyelectrolyte simulations (see the figure 3 in ref. 63). We invite the reader to watch the movie provided as SI to observe the formation of such a structure. We term this specific case of counterion bridge *nP-collab* (for *n* phosphate(s) collaboration) and define it as the presence of a bulk cation within 4 Å of *n* phosphorus atom from *n* different phosphate groups. This phenomenon was monitored along the trajectories, and we can see in Figures 3 and 4 that its propensity varies widely depending on the chosen force-field/water model combination. More precisely, simulations using the A19 and A99 systems present very few nP-collabs involving two phosphorylated serines (2P-collabs, in dark blue on Figure 3) and no nP-collabs involving three phosphorylated serines (3P-collabs, in red on Figure 3), while the laters are very common along the trajectories obtained with the A18 system. C36 simulations lead to an intermediate situation, with one replica (out of three) presenting nP-collabs during the whole trajectory and the two others having only transient nP-collab states.

**Figure 2:**
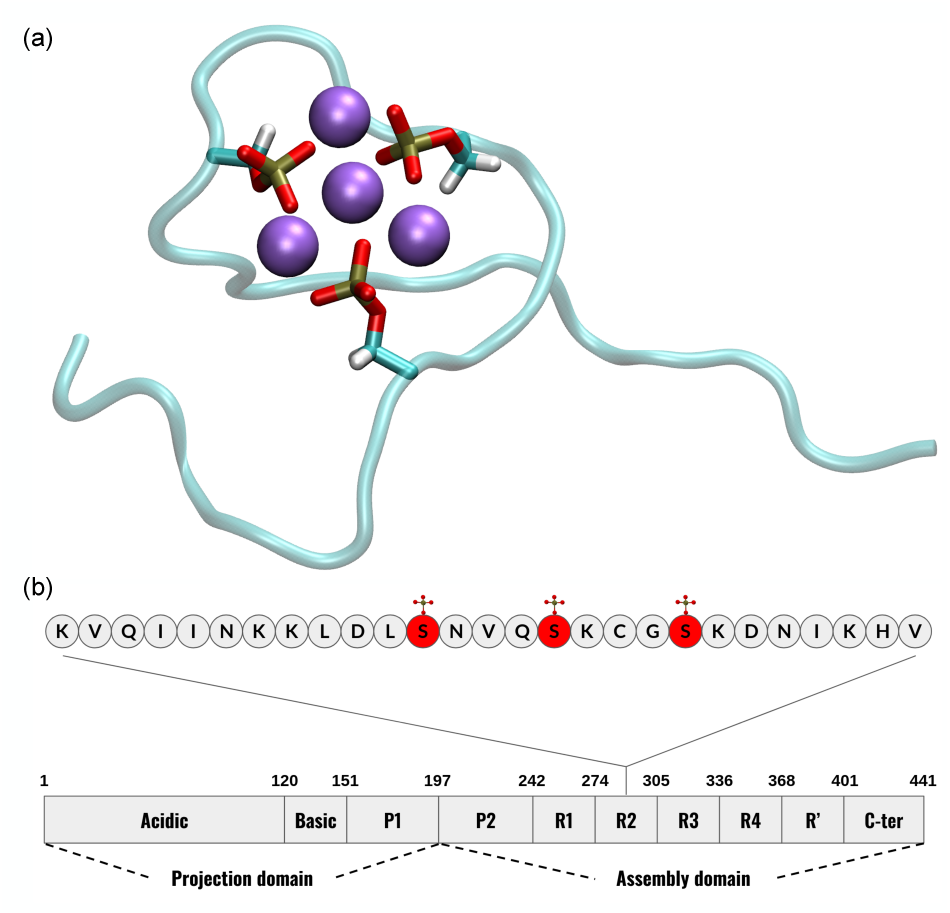
(a) Snapshot from the MD simulation (replica 1) of triply phosphorylated tau-R2 with the C36 system illustrating a 3P-collab situation with Na^+^ ions shown as purple van der Waals spheres. (b) Schematic diagram of tau protein highlighting the investigated phosphorylation sites in the R2-fragment.

**Figure 3:**
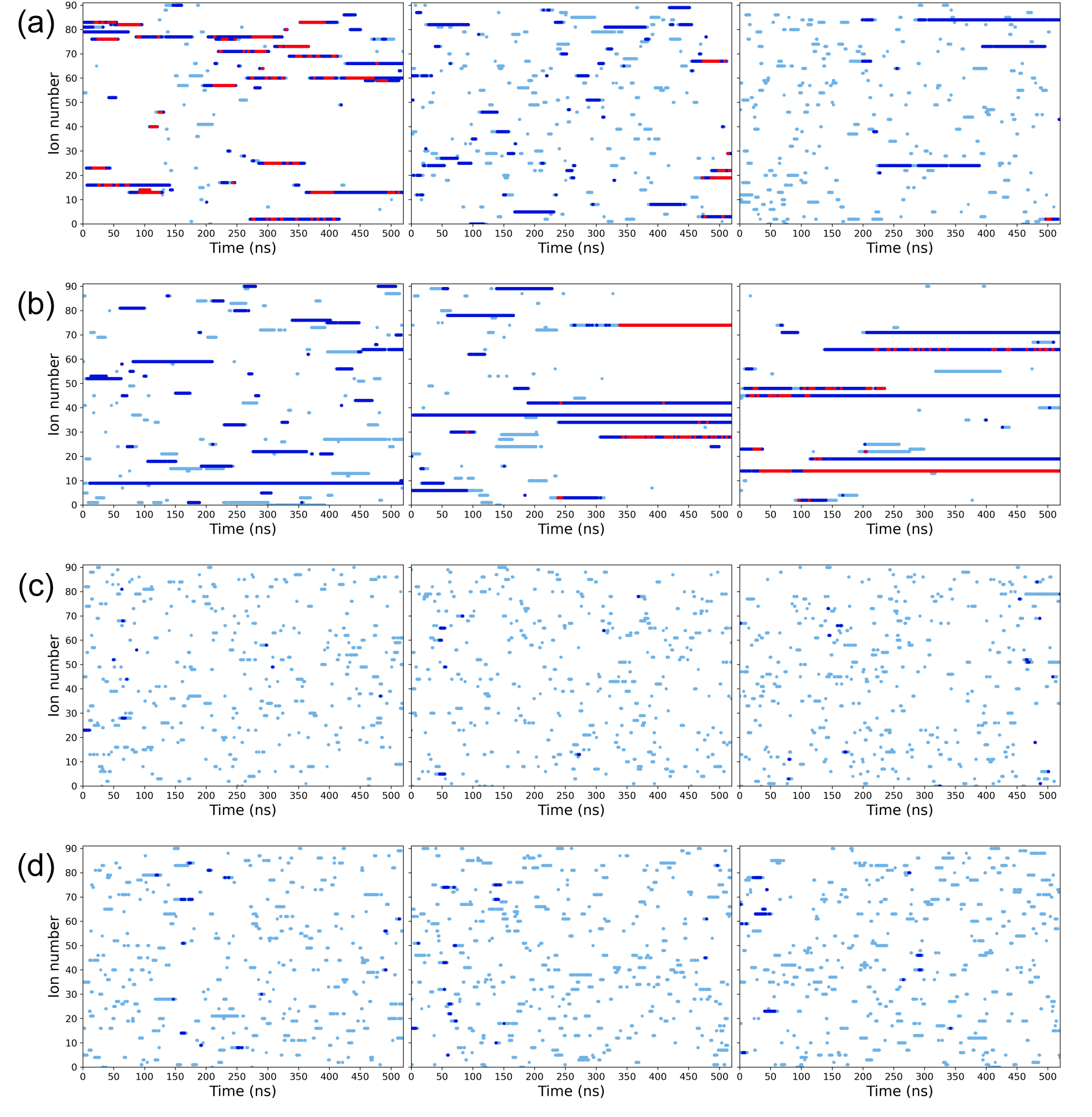
Monitoring the presence of cations in the vicinity of the phosphate groups along the MD trajectories with Na^+^ counterions for each one of three replicas. Each horizontal line corresponds to one individual cation. Light blue dots indicate that the cation is less than 4 Å away from a P atom, dark blue indicate proximity to two P atoms (2P-collabs) and red proximity to the three P atoms (3P-collabs). Force-field/water combination : **(a)** C36, A18, **(c)** A19, **(d)** A99

**Figure 4:**
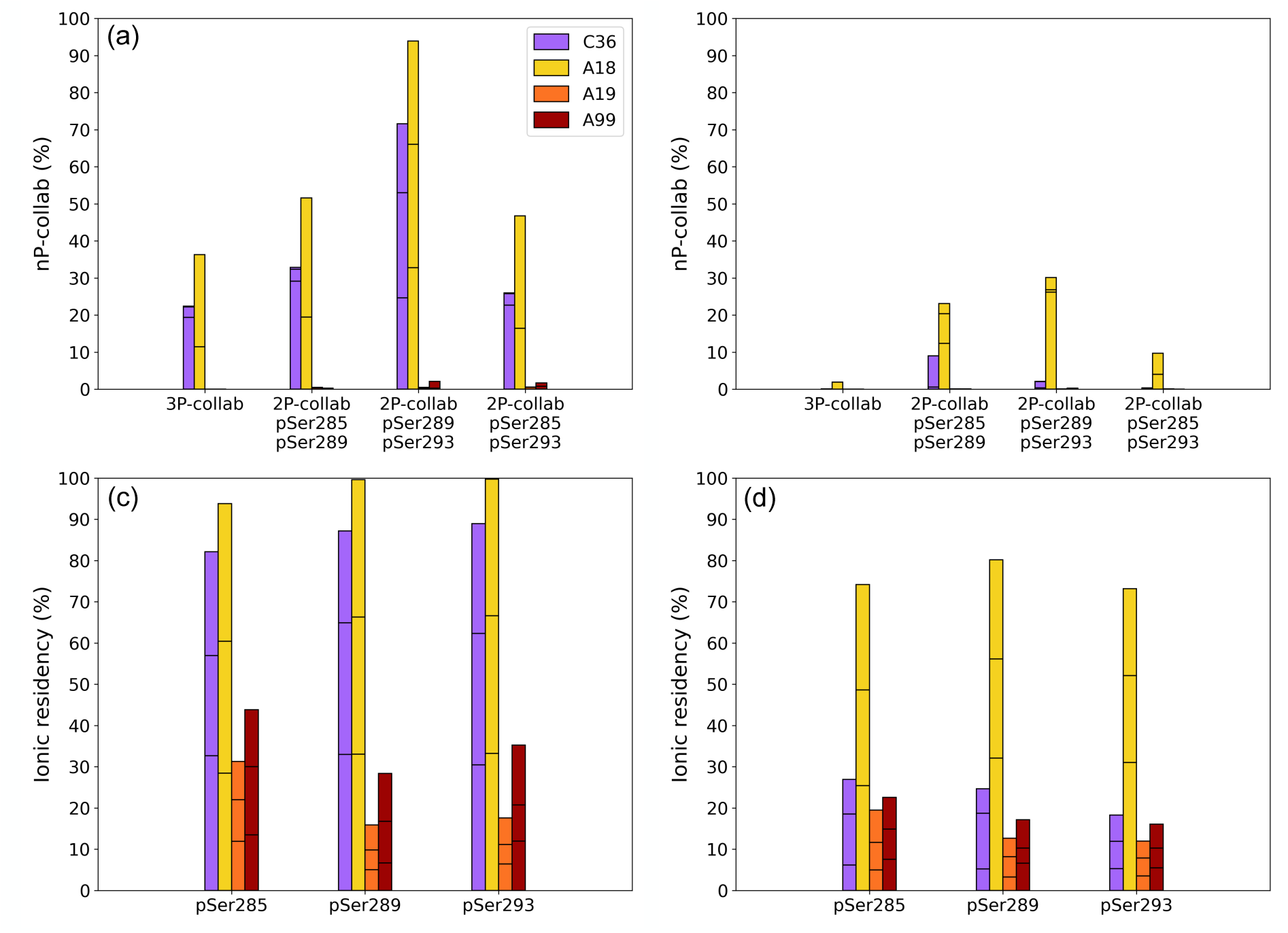
nP-collab (with a counterion located less than 4 Å away from each P atom) rates during the MD simulations over the three replicas (the segments in each column show the contribution of each replica to the global average). **(a)** Simulations with Na^+^ counterions, **(b)** Simulations with K^+^ counterions. Ionic residency rate (with a counterion located less than 4 Å away from the P atom) for each pSer residue in the phosphorylated R2-fragment. **(c)** Simulations with Na^+^ counterions, **(d)** Simulations with K^+^ counterions.

Another interesting feature of nP-collabs is that a phosphocollaborated cation is able to exchange its place with another cation from the bulk without compromising the inter-phosphate bridge. This behavior seems to be more likely in C36 than in A18 (see Figure 3a-b), making the former once again an intermediate situation. nP-collabs also tend to bend the structure of the peptide, increasing its LC (central panel in Figures 1b and 1d) and lowering the LF of some residues (right panel in Figures 1b and 1d). The LF metric allows a quantification of the complex impact of nP-collabs on the peptide dynamics. C36 and A18 are the only systems engaging in 3P-collabs, and a common consequence is a lowered LF between pSer289 and pSer293. Finally, the mobility of serines is reduced for all systems upon phosphorylation except for pSer293 in C36 due to the making and breaking of 3P-collabs.

### Cation-phosphate interactions are force-field specific

Changing from sodium to potassium counterions leads to a strong decrease of the nP-collab rate for all four systems, with nP-collabs now appearing almost only when using the A18 FF/ water model combination (see Figures 4b and SI-6). Moreover the sodium ionic residency (% of time a sodium ion spends within 4 Å of the phosphorus atom of a specific phosphorylated serine) is above 80% for C36 and A18, whereas it remains below 50% for A19 and A99 (Figure 4c). C36 displays the most striking decrease when changing from Na^+^ to K^+^ counterions: K^+^ is nearly three times less present around phosphoserines than Na^+^ (Figure 4d). We can note that this decrease in the counterion/phosphate interaction when comparing Na^+^ and K^+^ concurs with experimental and theoretical data in an early work of Schneider et al. surveying crystal structures.^64^

### Impact of the number of phosphorylations in tau-R2

For the C36 system, we also ran simulations with only one or two phosphorylated serines (see Table 1). As the formation of nP-collabs requires at least two phosphate groups, we investigated whether the position of these phosphorylated groups along the peptide sequence could impact the formation of such counterion bridges. Indeed, the 2P-collab rate between pSer285 and pSer293 is always lower than between neighboring phosphorylated serine pairs (pSer285-pSer289 or pSer289-pSer293), both when modeling a triply phosphorylated R2 fragment (Figure 4a-b), or a R2 fragment with only two phosphorylated serines (Figure SI-7).

### Sequence and concentration effects: 2P-collabs in a polyalanine toy model

In line with the approach of He and al.,^65^ in order to further investigate the impact of the spacing of the phosphorylated serines along the sequence on the nP-collab rate, we also performed MD simulations with the C36 FF/water model combination (and using the same protocol as described in the *Material and Methods* section) on a 13 residue-long polyalanine peptide comprising two phosphorylated serines in position i and i+j, with j varying from one till six (see Figure 5a). The initial structures for the toy models were built with the peptide builder of the tleap program available in the AmberTools package,^66^ and serines were phosphorylated using CHARMM-GUI. One should note that in this system, the phosphate groups cannot form salt-bridges with charged residues, meaning that the compaction of the peptide observed upon phosphorylation (see Figure SI-8) can only be attributed to the formation of 2P-collabs. These were observed for each configuration, as shown on Figure 5b.

**Figure 5:**
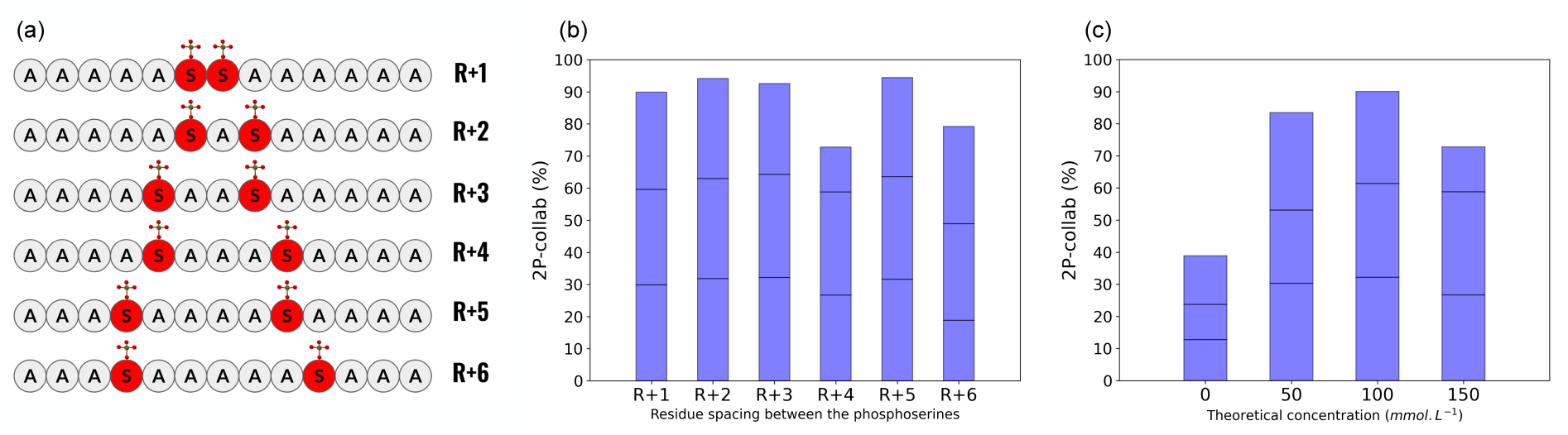
**(a)** Schematic of the polyalanine toy models with serines spaced in various positions **(b)** Evolution of the 2P-collab rate in the polyalanine toy models when varying the pSers spacing. **(c)** Evolution of the 2P-collab rate in the R+4 system when varying the ionic concentration. The 0 concentration corresponds to a system that comprises only four neutralizing cations. For panels b and c, the segments in each blue column show the contribution of each replica to the global average.

Local Curvature and Local Flexibility also reveal a spacing-dependent effect on the conformational variability and dynamics of the toy models. We used a peptide constituted solely of alanines as a reference. As can be observed on Figure 6, the behavior of the peptide changes when the spacing between the pSerines goes beyond 3 residues. For the R+1 to R+3 peptides, LC between the phosphoresidues is either unchanged or slightly reduced compared to the polyalanine reference state, whereas for the R+4 to R+6 peptides LC is drastically increased. This can be explained by the apparition of a hairpin in the backbone (see the snapshots in Figure 6) which progressively bends it. LF is reduced for the residues located between the phosphorylated serines for all peptides, indicating that 2P-collabs can be responsible for a rigidification of the backbone. LF also hints at a peculiar behavior of the peptide R+6 as it is the only toy model presenting a drastic increase of flexibility around a pSerine (see Figure 6f). Looking at the PMC for each replica of peptide R+6 allows to identify two distinct behaviors of the peptide (see Figure SI-9). The 2P-collab can either create a sharp hairpin centered on Ala7 (Figure SI-9, central and right panels) or allows the creation of a flattened hairpin with a short α-helix involving the five N-ter residues (Figure SI-9, left panel).

**Figure 6:**
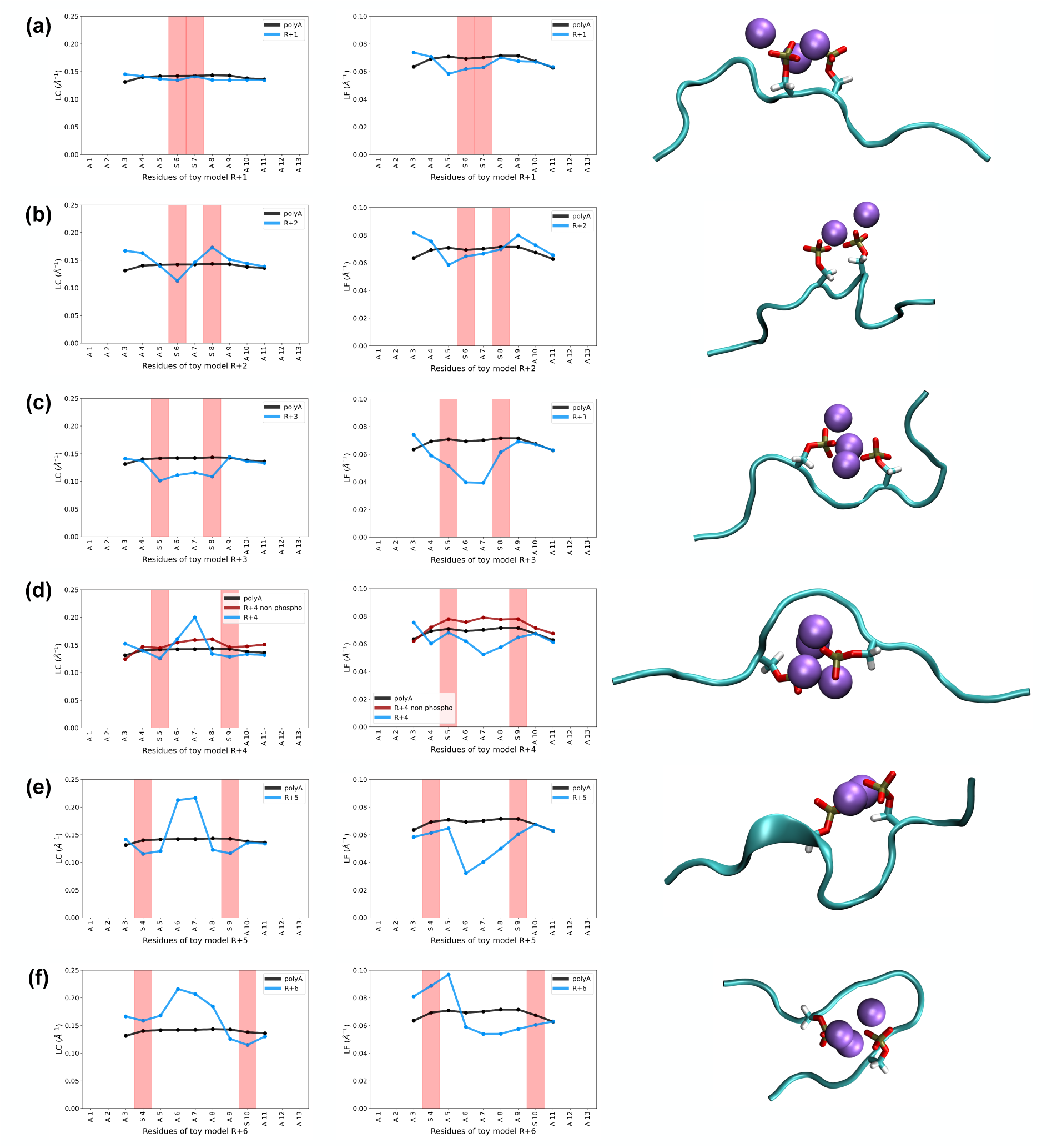
Local curvature (LC, left panel) and flexibility (LF, central panel) in the polyalanine toy models with representative snapshots (right panel) of the conformational states with 2P-collabs. Black line : reference polyalanine peptide, blue line : phosphorylated peptide. The red columns in the left and central panel figures highlight the position of the phosphorylated serines along the sequence. **(a)** R+1 model, **(b)** R+2 model, **(c)** R+3 model, **(d)** R+4 model, the additional red line corresponds to the peptide with unphosphorylated serines, **(e)** R+5 model, **(f)** R+6 model.

For the system with pSerines placed four residues apart, we also investigated the impact of the ionic force. Interestingly, using counterions only to neutralize the system (which corresponds to only four Na^+^ or K^+^ cations in the simulation box and to the concentration of 0 on Figure 5c) is sufficient to observe an average 2P-collab rate of 40% over the three trajectories. We can also observe the impact of phosphorylations on the tau-R2 conformations by looking at the Ramachandran plots resulting from the MD trajectories. Figure 7 shows the evolution of the ϕ/ψ angles of the Ala6 residue in the R+4 toy model, which were sensitive to the introduction of serines in positions 5 and 9 and their following phosphorylation.

**Figure 7:**
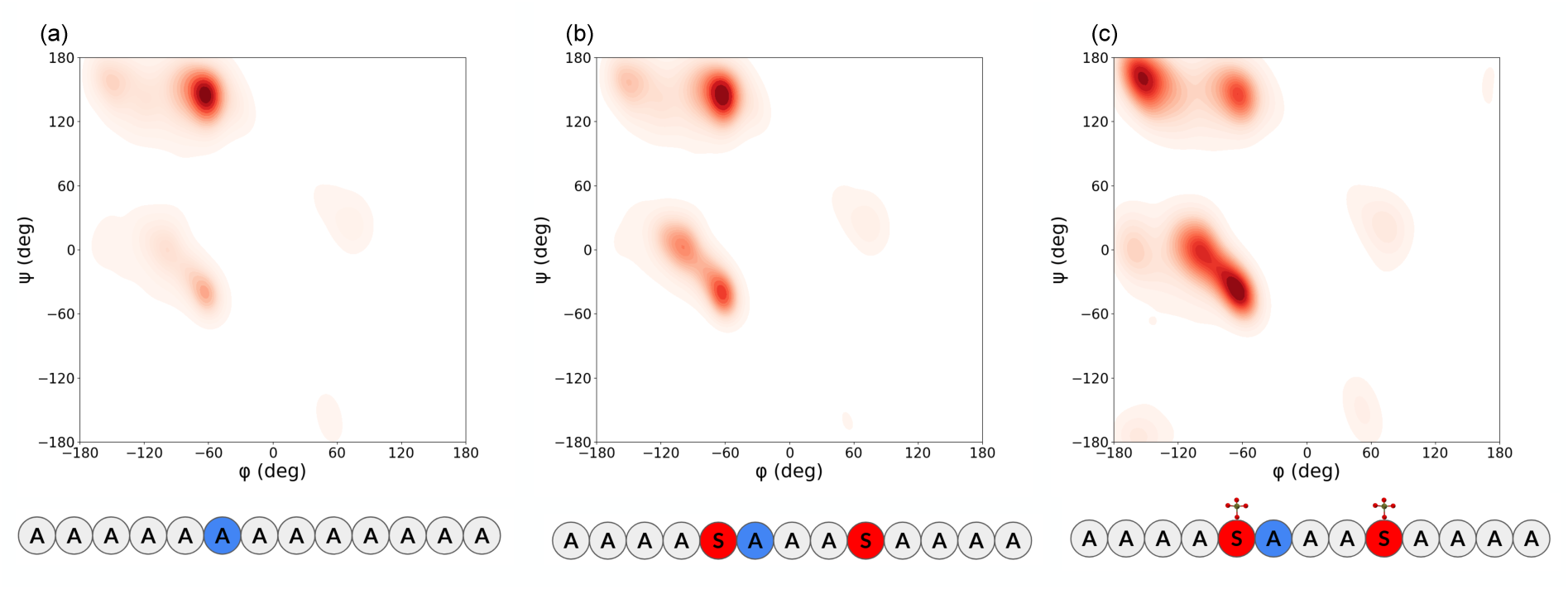
Ramachandran plots for Ala6 in the R+4 toy model **(a)** polyalanine toy model **(b)** toy model with serines in positions 5 and 9 **(c)** toy model with phosphorylated serines in positions 5 and 9.

### Parametrization of the phosphate groups in the different force-fields

The behavior difference observed between the four FF/water combinations can first be related to the parametrization of the phosphate groups, and more specifically their rigidity. Figure 8 shows the distribution of the three dihedral angles χ_1_, χ_2_ and χ_3_ (defined in Figure 8a) during the MD simulations, and we can see how the χ_2_ angle is much more rigid for the A19 and A99 FFs, thus preventing the reorientation of the phosphate group to form nP-collab structures. This is in agreement with the distributions observed by Vymetal et al. when comparing the structure of phosphorylated serines with the A99 and C36 FFs.^67^ A19 also displays very narrow distributions of χ_1_ and χ_3_ . This was more unexpected, as the phosaa19sb phosphate parameters used in the A19 system derive from the phosaa10 parameter of the A99 system, and in their paper on these phosaa10 parameters, Homeyer et al. state that “For the phosphate groups of phospho-amino-acid residues, free rotation around the O-phosphomonoester or phosphoramidate bond would be expected”.^52^ Conversely, the flexible dihedral angle distributions obtained for the C36 and A18 systems agree with the experimental data obtained by Zang et al. in their review of PTMs in the PDB.^68^ In addition, the Lennard-Jones and Coulomb parameters for the counterions (Na^+^ or K^+^) interacting with oxygen from water or from phosphate groups appear to be clearly in favor of water oxygen in the case of the A19 and A99 systems, while the C36 and A18 systems favor cation/phosphate interactions (see Figure SI-10). The theoretical non-bonded potentials also reveal that the counterions/ phosphate oxygens interaction is the least favorable for A19, which is coherent with it being the least likely to form nP-collabs (see Figures 4a-b). We thus argue that, from a strictly computational point of view, nP-collabs require two necessary conditions to occur: that the rigidity of the phosphoserine sidechain is low enough to allow reorientations, and that the non-bonded parameters of the force-field induce bulk cations to be more attracted by terminal side-chain phosphate oxygens than by water oxygens.

**Figure 8:**
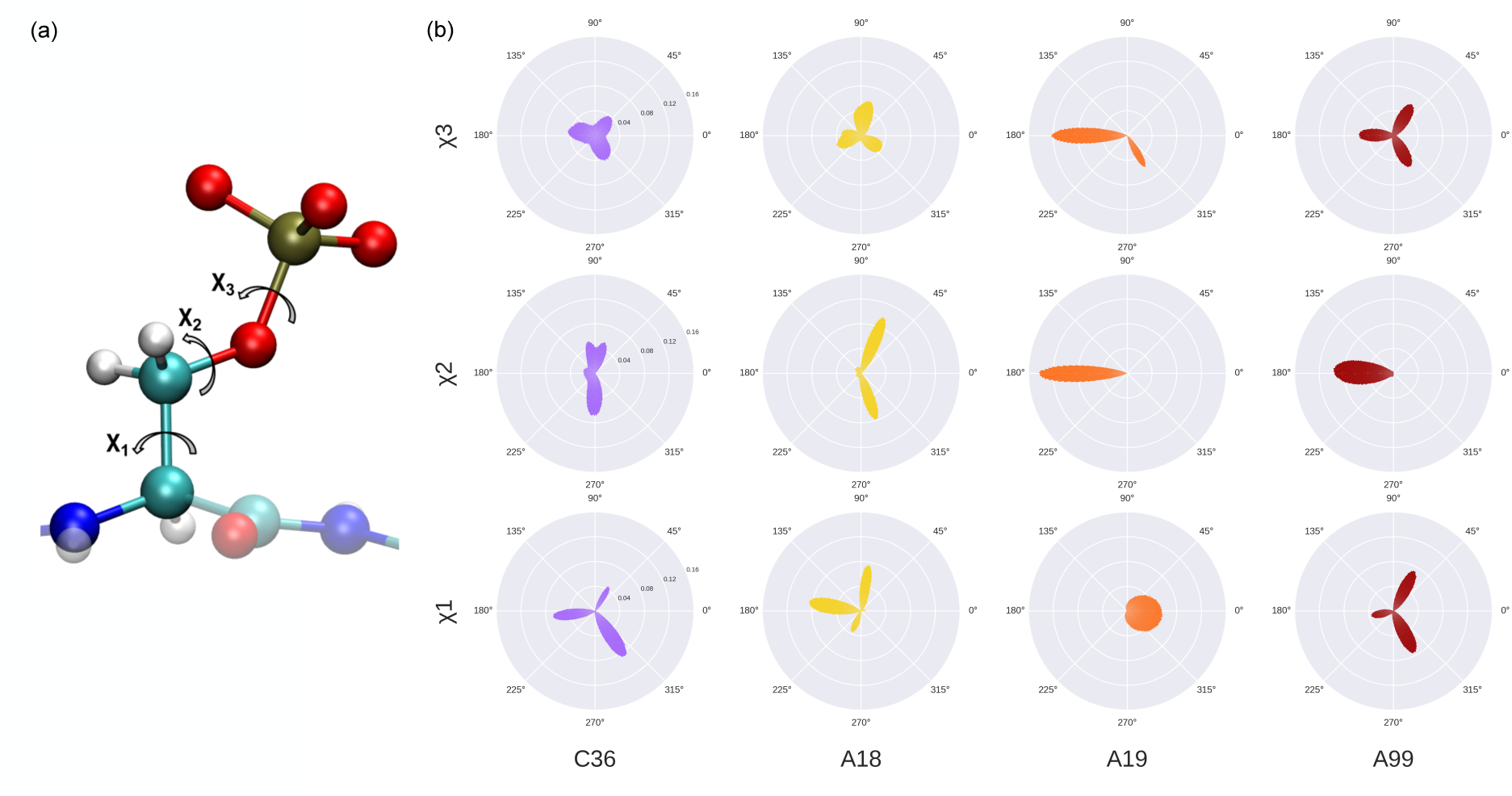
(a) Dihedral angles χ_1_, χ_2_ and χ_3_ in a phosphate group. **(b)** Distribution of the dihedral angles in pSer289 over the trajectories of the three replica for the triply phosphorylated system in four different FFs.

## Discussion

### Earlier observations of P-collabs

Most theoretical works dealing with the phosphorylation of IDPs focus on the formation of salt bridges involving positively charged residues (mainly lysine and arginine), without explicitly investigating the role played by counterions in the system. However, we found several works that agree qualitatively with our findings, in particular when comparing different force fields. For example, Rieloff and Skepö performed a series of studies on IDPs with and without phosphorylations comparing the A99 and C36 FFs,^29, 69^ with C36 leading to more compact conformational ensembles than A99, and several snapshots showing neighboring phosphate groups facing each other hint at possible 2P-collabs (see for example figure 6 and 7 in ref. 69). Recent work by Song et al. with FB18 (and sodium counterions for system neutralization only) clearly displays 2P-collabs in the 2CEF PDB structure, contributing for 64.1% of the total conformational ensemble (see figure 8 of ref. 70). Their improvement of FB18, FB18CMAP, also seems to lead to the formation of 2P-collabs, up to 43.5% of the conformational ensemble, which is more in line with the proportions obtained using C36. Amith and Dutagaci used MD simulations with the C36 FF to investigate the impact of multiple phosphorylations on the conformation space explored by the C-terminal domain of RNA polymerase II.^71^ While the study was performed in neutralizing conditions without added ions, the counterions tended to condense around the phosphate groups and nP- collabs involving up to four phosphorylated serines could be observed. 2P-collabs are present in the trajectories produced by Bickel and Vranken^72^ in their extensive study of mono- and diphosphorylated peptides using MD simulations with the A99 FF/water model combination and a NaCl concentration of 0.15 mol.L^-1^. In particular, 2P-collabs also formed in peptides with other phosphorylated residues such as phosphothreonine and phosphotyrosine.

Intermolecular 2P-collabs were also observed by Man et al. when modeling a system comprising two monophosphorylated (pSer289) tau-R2 fragments with the C36 FF.^73^ A pS289-Na^+^-pS289 bridge was present in 16% of the MD trajectory, and this result raises the question of whether nP-collabs could be a possible pathway to aggregation by bridging monomers. Generally speaking, all these recognized and unrecognized sightings of nP-collabs make a strong case for the importance of bulk ions in MD simulations, and we therefore urge the computational community to reconsider their relevance and keep track of them in the future. As classical all-atom forcefields are known to suffer from their lack of explicit description of dipole effects, polarizable forcefields such as AMOEBA^74^ or CHARMM- Drude^75^ could therefore provide a more accurate representation of the cation-phosphate interaction. Going even further down the fundamental lane, QM/MM simulations would be excellent candidates for assessing the stability of nP-collabs at the quantum level, but suffer from inherent limitations which make them difficult to implement with highly dynamic systems such as IDPs.^76^ Both polarizable forcefields and QM/MM simulations mostly suffer from their high computational cost, yielding timescales that might be too short to properly investigate this phenomenon. The Electronic Continuum Correction (ECC) scaling method is based on a mean field theory treating electronic polarization as a dielectric continuum,^77^ and could provide an interesting compromise between the costly polarizable forcefields and the dipoleless classical forcefields. Meanwhile, a temporary fix for classical forcefields could be the use of the NBFIX method, which introduce pair-specific nonbonded interaction parameters.^78^ As a matter of fact, this type of correction was already introduced between sodium cations and certain lipid phosphate oxygens in the CHARMM forcefield, providing a relevant proof of concept for future applications to corrections of nP-collab interactions.^79^

From the experimental perspective, Lasorsa et al. recently combined NMR spectroscopy and MD simulations using the C36 and A99 FFs to investigate the conformation space explored by another tau fragment (210-240) with multiple phosphorylations (on T212, T217, T231 and S235).^80^ Their comparison of experimental and theoretical data suggest that the experimental parameters are best captured by simulations using the CHARMM36m FF with TIP4P-D water with respect to the A99 simulations. Interestingly, they also mention that a single phosphorylation of the tau fragment might not be sufficient for stabilizing compact structures of the peptide, thus suggesting that nP-collabs, and not only the salt bridges formed between phosphates and positively charged residues, might play a part in the compaction phenomenon observed upon phosphorylation. In another recent work, Lasorsa et al. performed magnetic resonance experiments on the AT8 epitope of tau, which was phosphorylated on Ser202 and Thr205.^81^ Their analysis of the relaxation rates of the NMR signal suggests that phosphorylation decreases the protein local dynamics around the phosphorylation sites which concurs with the flexibility plot we obtained for the polyalanine model with a R+3 phosphate spacing (see the central panel in Figure 6c). Finally, we used AlphaFold3^82^ to predict the structure of the triply phosphorylated tau-R2 repeat, and the resulting predictions do include nP-collabs (see Figure SI-11), thus supporting the idea that such structures are indeed present in the experimental data from the PDB used to train the model. However, experimentally detecting something as elusive as the dynamic interactions between monovalent cations and phosphate groups on IDPs might prove a remarkable challenge. Our guess is that it could be easier to detect the conformational rigidification of the backbone induced by possible nP- collabs. We already discussed the insights provided by magnetic resonance techniques such as NMR, and this class of experiments has shown great sensitivity to solvent-associated changes in IDP dynamics before.^83^ SAXS is another technique which proved to be valuable for deciphering IDP dynamics and its relation to outer conditions.^84^ We suggest that experiments could be performed on our toy models in order to directly compare results such as the radius of gyration or local flexibilities. One main concern we would like to raise is the possibility for nP-collabs to be inter-monomer, which could make the correlation between single-molecule simulations and in vitro concentrated experiments not straightforward.

## Concluding remarks

We ran classical Molecular Dynamics simulations to investigate the formation of counterions bridges, involving at least two phosphate groups and a cation and named nP-collabs, in the triply phosphorylated, disordered, R2 fragment of tau. Using four different FF/water-model combinations, we show that the formation of nP-collabs in tau-R2 during the simulations is highly dependent on the chosen simulation parameters. Further investigations of the phosphate group parametrization show that this phenomenon can be related to the degree of rigidity of the phosphate groups, and to a favorable cation/phosphate oxygen interaction compared to the cation/water oxygen one. The development of three new metrics, Proteic Menger Curvature, Local Curvature and Local Flexibility allowed us to fully characterize the behavior of our multiphosphorylated disordered toy models, and show promising results for general investigations of the conformational ensembles of IDPs and relatability to NMR measurements.

Searching the scientific literature showed that nP-collabs were already observed on several occasions in earlier computational studies dealing with various cases of phosphorylated IDPs, while seldom being explicitly addressed, one reason being that the counterions are usually not included in the analysis of the trajectories and conformational ensembles produced by MD simulations. However, nP-collabs, in particular when working with sodium counterions, are very likely to play a part in the structural and dynamic changes observed in IDPs undergoing multiple phosphorylations, and they might also impact their binding properties. Several experimental results hinting to the possible existence of nP-collabs were also discussed, but further in-vitro investigation will have to be conducted in order to confirm their existence. Indeed, the pseudo-structuration induced by nP-collabs could also provide new insights for experimental approaches or even drug design, tauopathies such as Alzheimer’s disease being of particular relevance.

In this perspective, future work will address the impact of tau-R2 phosphorylation on its interaction with the tubulin surface. In particular, we plan to pay specific attention to the formation (or not) of nP-collabs in tubulin-bound tau-R2 upon multiple phosphorylation, and how this phenomenon could change the way tau-R2 binds on the microtubule surface.

## Supporting information

AlphaFold3 structural prediction for the phosphorylated tau-R2 fragment

Movie showing a 3P-collab formation in the phosphorylated tau-R2 fragment

## Acknowledgments

The authors would like to thank J. Puyo-Fourtine and É. Duboué-Dijon for fruitful discussions regarding the parametrization of ions in classical all-atom FFs. The authors would like to thank D. Bickel and W. Vranken for providing us with additional trajectories from phosphorylated peptides to perform our analyses. This work was supported by the ANR (MAGNETAU-ANR-21-CE29-0024) and the «Initiative d’Excellence» program from the French State (Grant « DYNAMO », ANR-11-LABX-011-01 and grant "CACSICE", ANR-11- EQPX-0008). Calculations were performed using HPC resources from GENCI-TGCC (grant A0140714126) and from LBT-HPC.

## Supporting Information Available

Additional data regarding the simulation parameters in the four considered force field/water combination, discussions regarding the ionic mobility and the convergence of the simulations, the tau-R2 contact maps and secondary structures along the MD trajectories, and when using K^+^ counterions.

A movie of a trajectory showing the formation of a 3P-collab in the phosphorylated Tau-R2 fragment.

An archive (.zip) with the structural predictions of the AlphaFold server on the triply phosphorylated tau-R2 fragment.

## Data and Software Availability

Python scripts developed for the calculation of the curvature metrics are available in a public G i t H u b r e p o s i t o r y : https://github.com/Jules-Marien/Articles/tree/main/_2024_Marien_nPcollabs

All discussed MD trajectories, without water (but including ions), and their topologies are deposited in Zenodo: https://zenodo.org/records/12634115

The Zenodo repository also includes files regarding the convergence of the simulations, DSSP plots for each trajectory discussed in the manuscript, and Local Flexibility plots for each replica for each simulation discussed in the manuscript.

## Notes

https://github.com/Jules-Marien/Articles/tree/main/_2024_Marien_nPcollabs

https://zenodo.org/records/12634115

### Competing Interest Statement

The authors have declared no competing interest.

### Summary of Updates

Additional discussions regarding the simulation convergence in the SI file and extended discussion on polarizable FFs use and possible experimental setups in the discussion section.

