## Supplementary information for "nP-collabs: Investigating counterion mediated bridges in the multiply phosphorylated tau-R2 repeat"

*Jules Marien[1], Chantal Prevost[1] and Sophie Sacquin-Mora[1]**

[1]Université Paris-Cité, CNRS, Laboratoire de Biochimie Théorique, 13 rue Pierre et Marie Curie, 75005, Paris, France

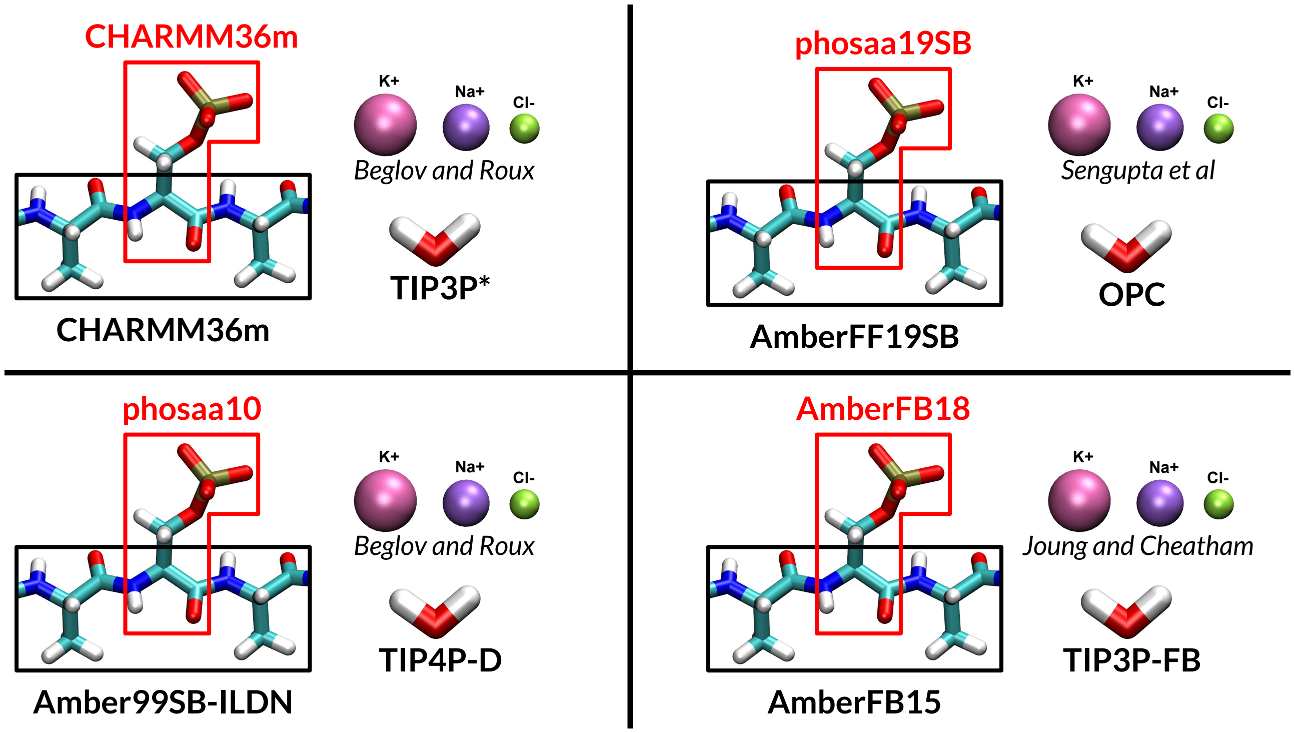


**Figure SI-1:** Visual summary of the force-fields/water models/ionic parameters/phosphate parameters used in the manuscript.

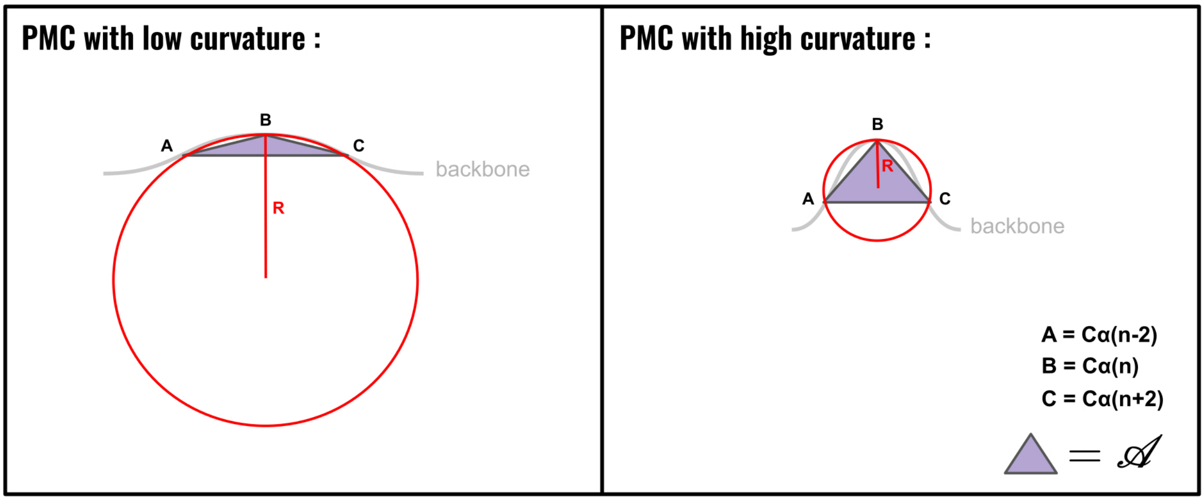


$$PMC = \frac{1}{R} = \frac{4\mathscr{A}}{AB \times BC \times AC}$$

$$LC = \langle PMC \rangle_c$$

$$LF = \sigma(PMC)_c$$

**Figure SI-2:** Visual summary for the Proteic Menger Curvature (PMC), the Local Curvature (LC) and the Local Flexibility (LF) for a given peptide conformation *c*.

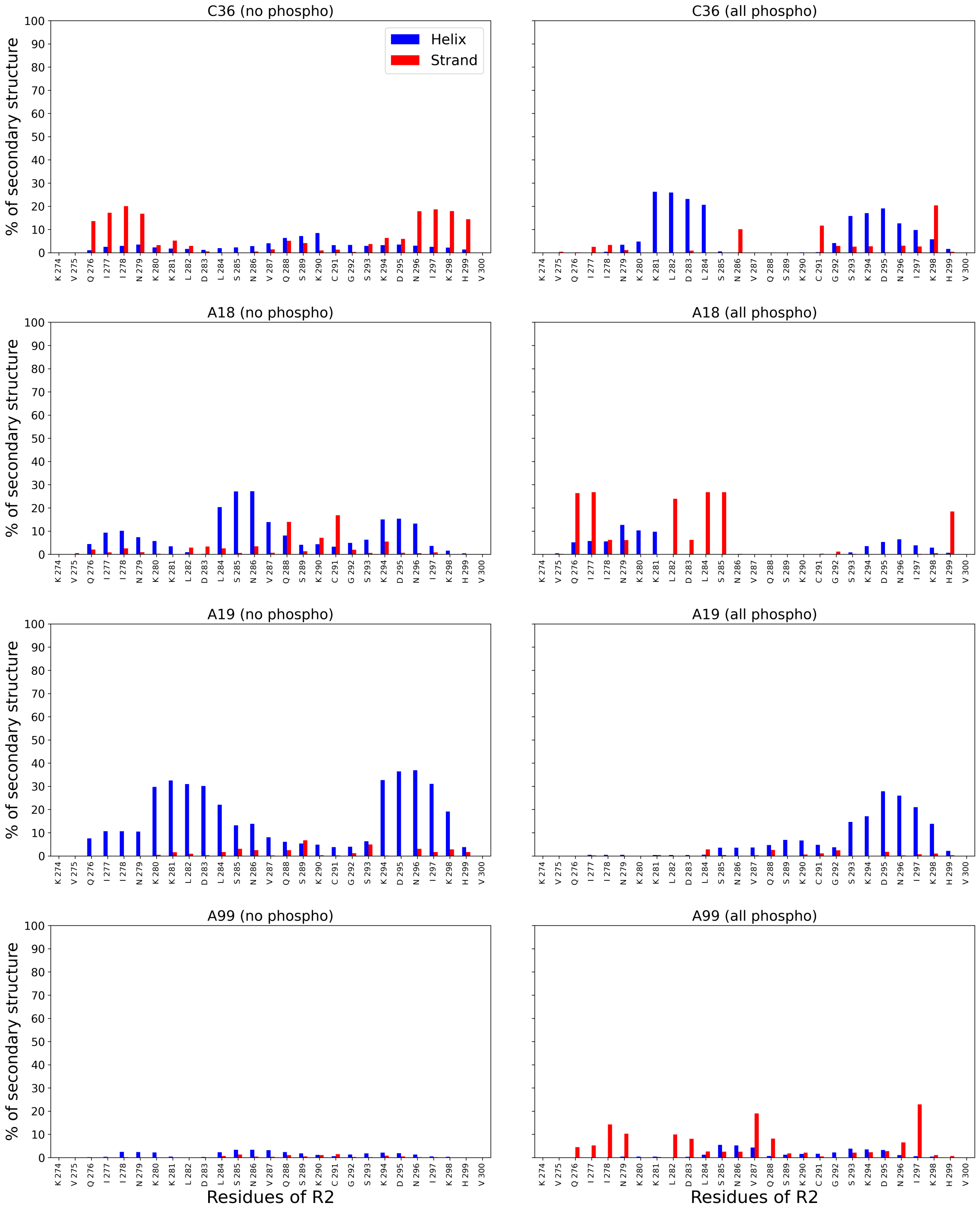


**Figure SI-3:** Helical (in blue) and β-sheet (in red) propensities of the tau-R2 residues for each force field in the unphosphorylated and triply phosphorylated states. Simulations with $Na^+$ counterions.

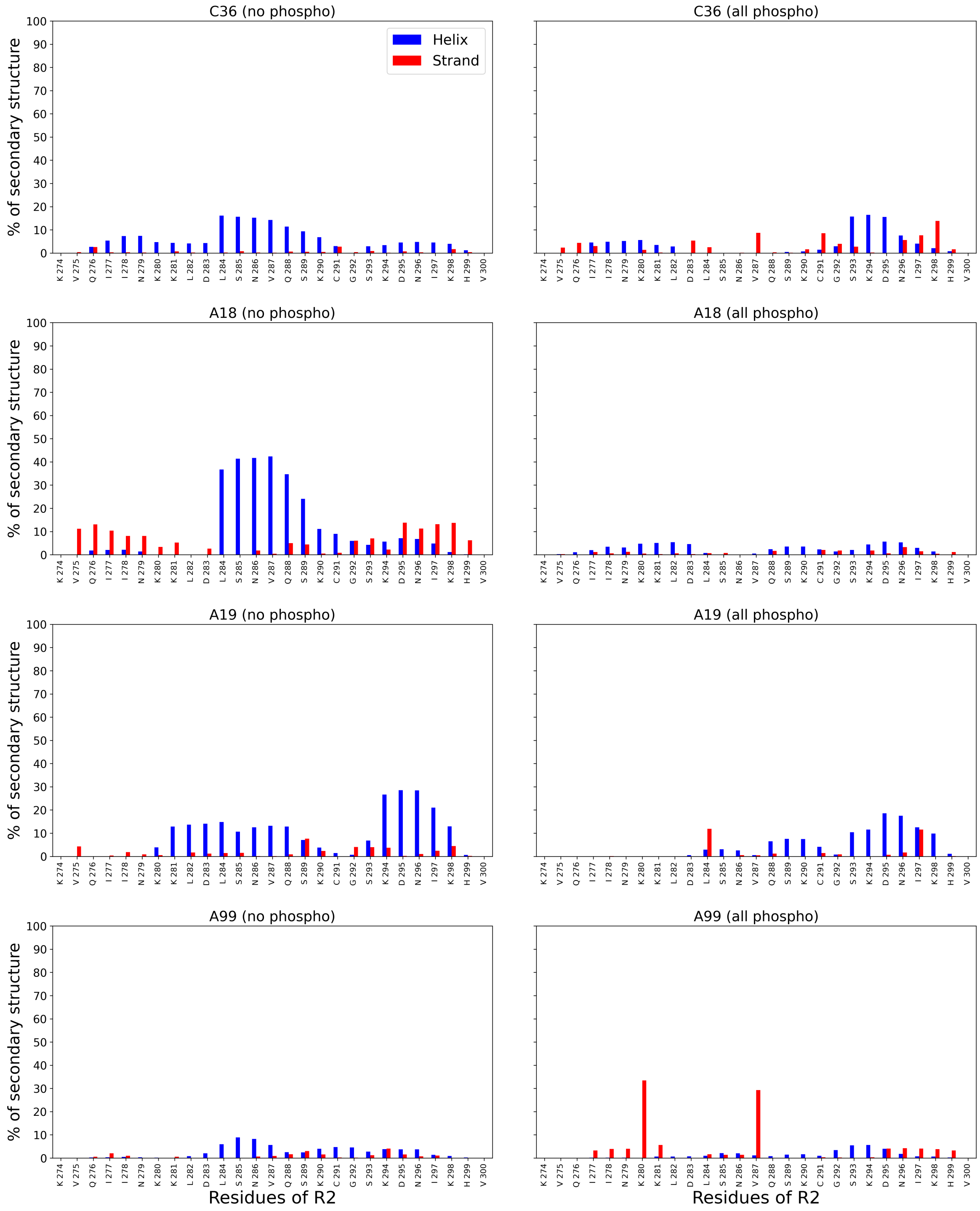


**Figure SI-4:** Helical (in blue) and β-sheet (in red) propensities of the tau-R2 residues for each FF/ water model in the unphosphorylated and triply phosphorylated states. Simulations with $K^+$ counterions.

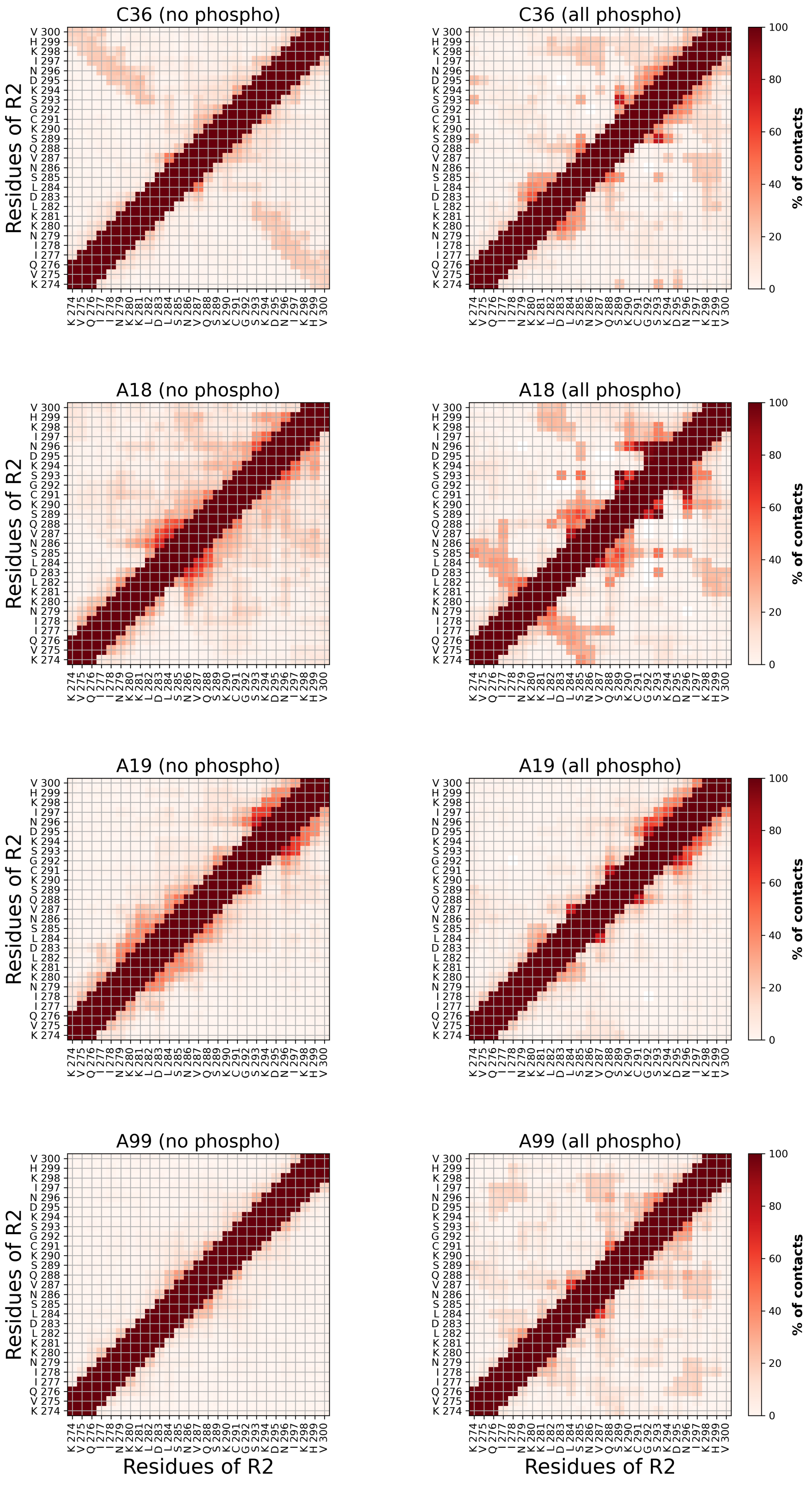


**Figure SI-5:** Contact maps between the tau-R2 residues (averaged over the three replicas for each system) with $Na^+$ ions, for the four different FF/water models, without and with phosphorylations.

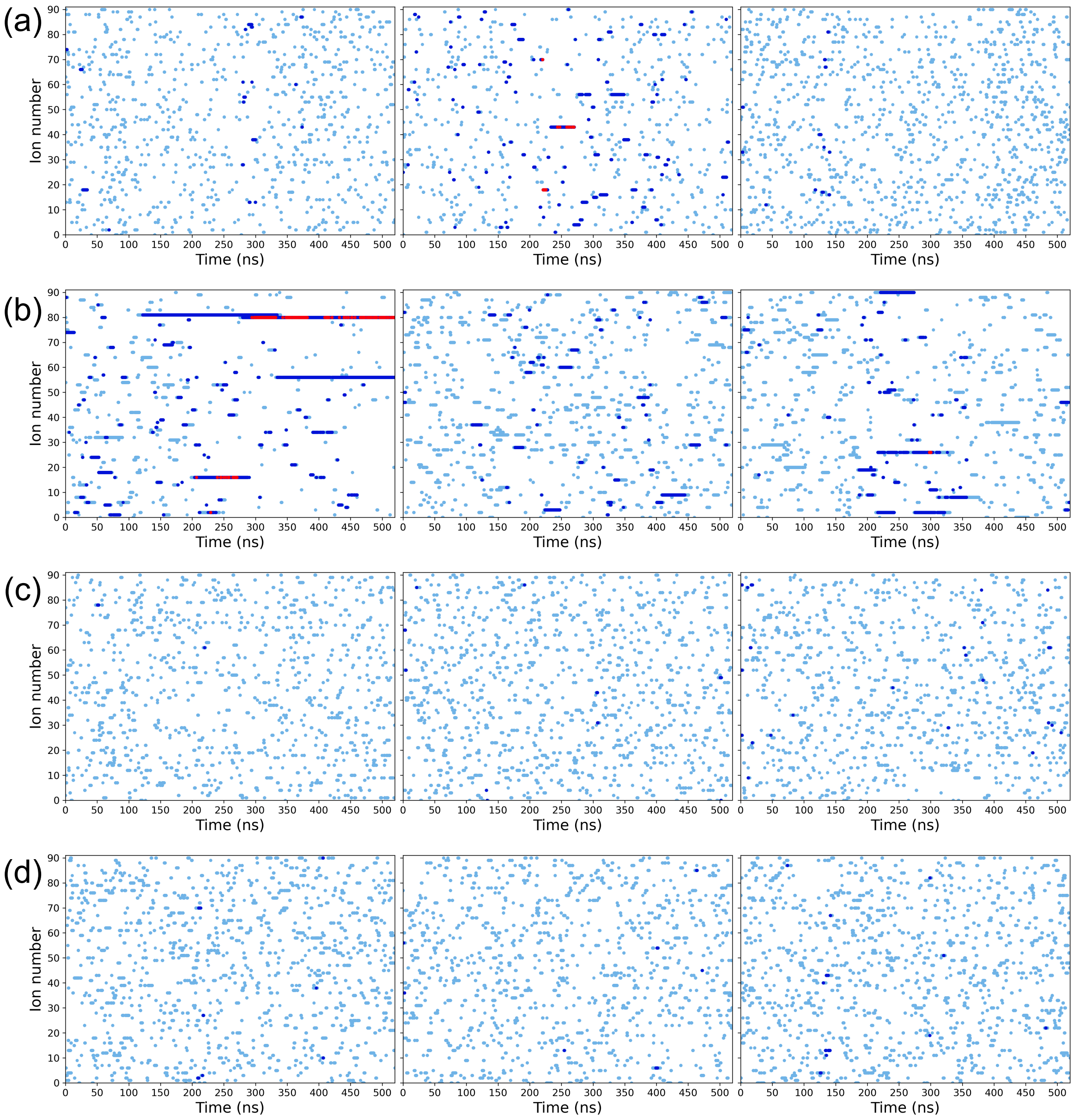


**Figure SI-6:** Monitoring the presence of cations in the vicinity of the phosphate groups along the MD trajectories with $K^+$ counterions for each one of three replicas. Light blue dots indicate that the cation is less than 4 Å away from a P atom, dark blue indicate proximity to two P atoms (2 serines P-collab) and red proximity to the three P atoms (3 serines P-collab).

Force-field/water model combination : **(a)** C36, **(b)** A18, **(c)** A19, **(d)** A99

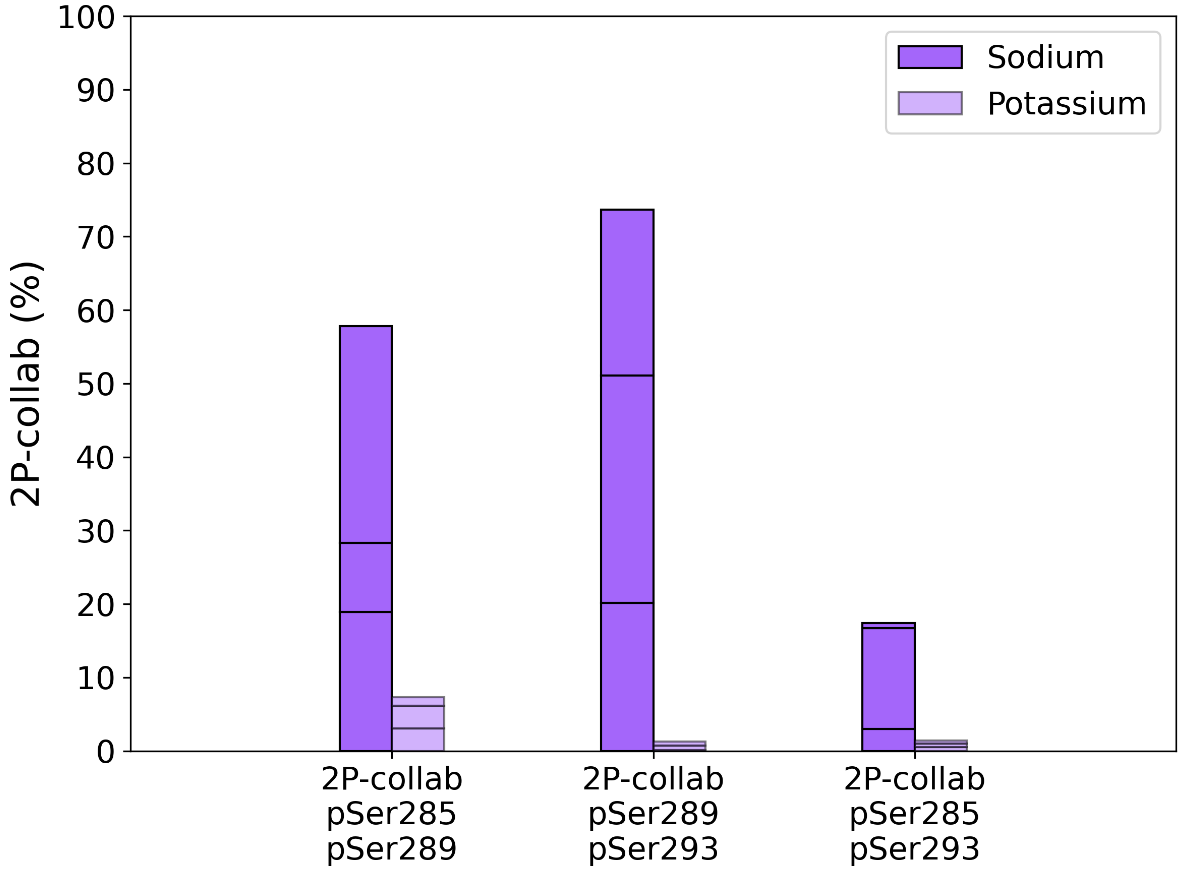


**Figure SI-7:** 2P-collab rate in tau-R2 with two phosphorylated serines modeled with the C36 system. The segments in each column show the contribution of each replica to the global average value.

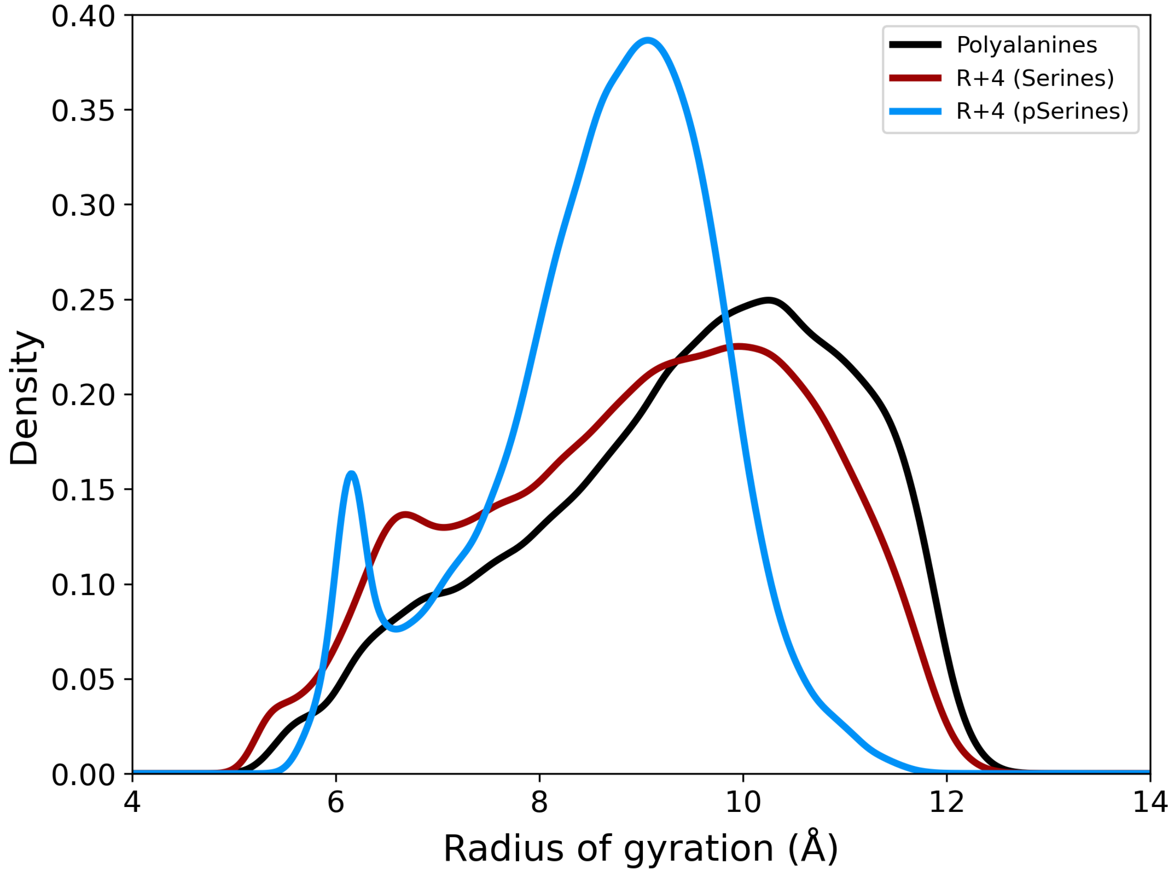


**Figure SI-8:** Density distribution of the radius of gyration of the polyalanine toy model (black), upon adding serines in positions 5-9 (dark red), after phosphorylation of the 5-9 serines (blue) with the C36 system.

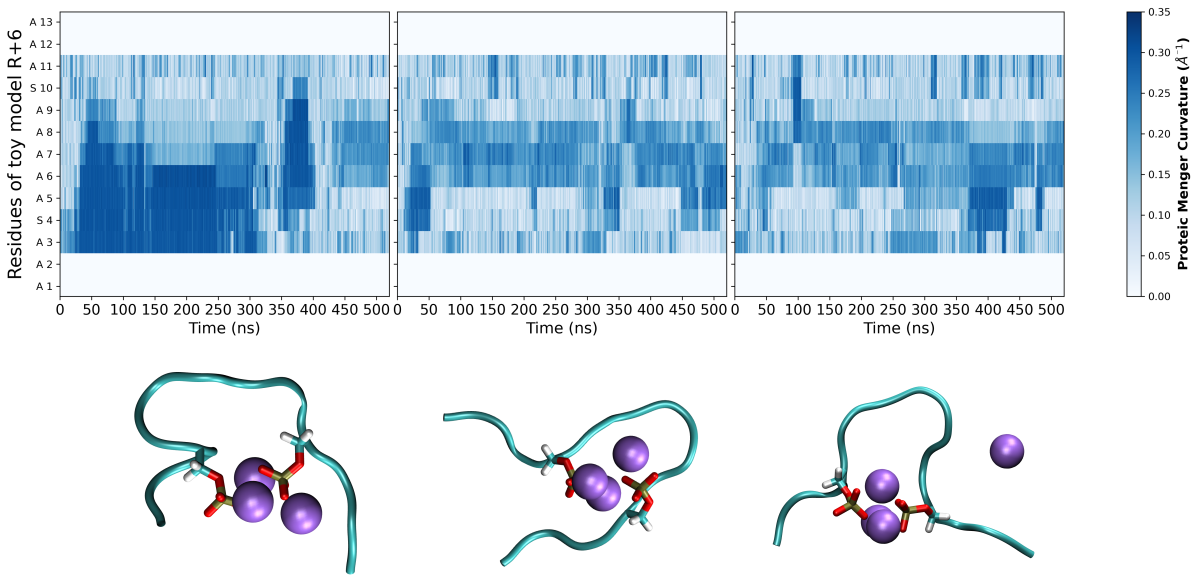


**Figure SI-9:** Top: PMC for each residue as a function of time in the R+6 toy model for the three replica. Bottom: Representative snapshots from each replica showing a 2P-collab situation with $Na^+$ ions shown as purple van der Waals spheres.

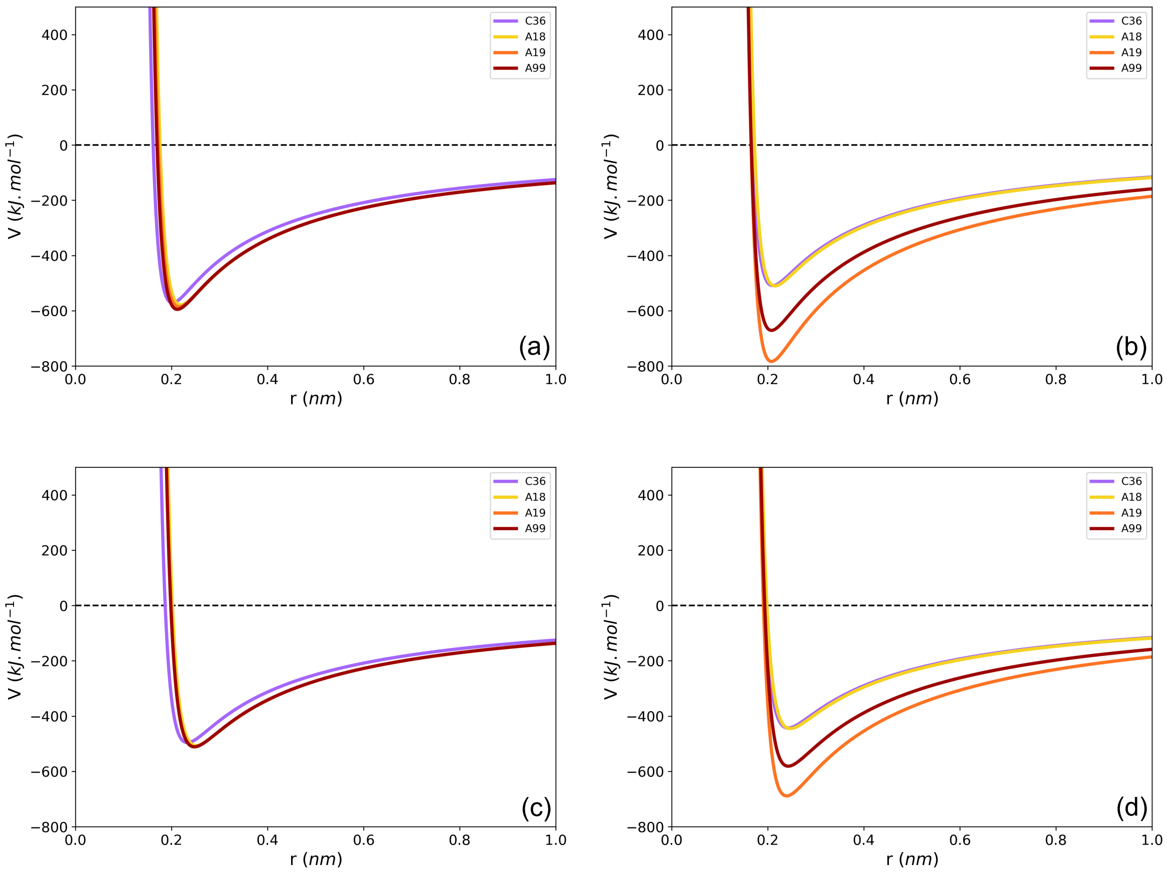


**Figure SI-10:** Cation/oxygen interaction as a function of the distance **(a)** $Na^+$ and phosphate oxygen, **(b)** $Na^+$ and water oxygen, **(c)** $K^+$ and phosphate oxygen, **(d)** $K^+$ and water oxygen.

The interaction potential is the sum of the Lennard-Jones and the Coulomb potentials. The potential analytical equations and parameters are available in the Github repository (https://github.com/Jules-Marien/Articles/tree/main/_2024_Marien_nPcollabs).

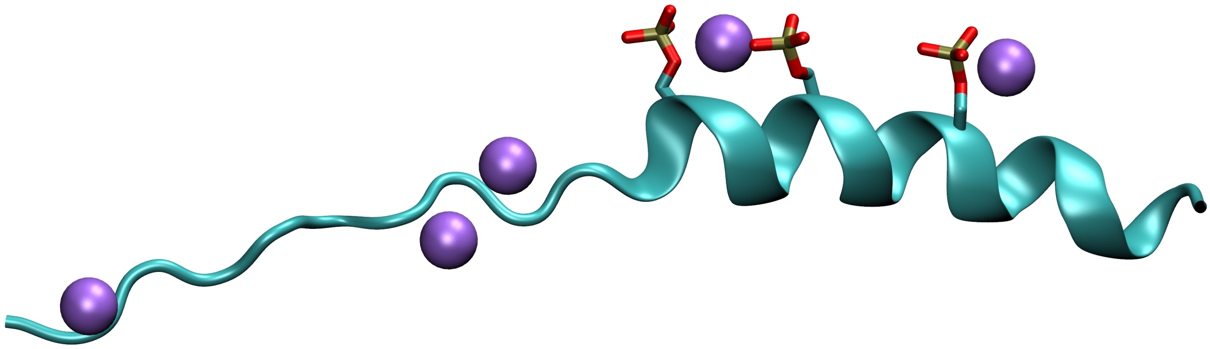

**Figure SI-11:** Structure prediction of the triply phosphorylated tau-R2 fragment with  5 $Na^{+}$ ions by AlphaFold3. Note that the helix structure on the C-terminal side is characteristic of the AF tools failure to treat disordered systems.

### Discussion on the simulation parameters

CHARMM36m benefits from a clear set of instructions regarding its use with GROMACS, available at the following address : https://manual.gromacs.org/current/user-guide/force-fields.html. We built a box of sidelength 10nm with gmx_editconf before filling it with water using gmx_solvate. We then specified the number of anions ($Cl^-$) and cations ($Na^+$ or $K^+$) with the gmx_genion command to keep the number of ions constant between similar systems, basing ourselves on a test output with the -conc flag set to 0.15 mol. We then minimized the systems.

Transferring an Amber Forcefield to GROMACS is not as straightforward. We however designed a robust pipeline to do so :

1. Start with AmberTools22[1]

2. Load the system, add water and ions, and create the coordinates (.inpcrd) and topology (.prmtop) files using tleap

3. Transform the .inpcrd file into a .gro file and the .prmtop file into a .top file using the amb2gro_top_gro.py script available in AmberTools22

4. Minimize the system

For both CHARMM36m and the Amber forcefields, we set the threshold of the steepest descent minimization function to 10.0 $kJ.mol^{-1}.nm^{-1}$ (default value) but none of the systems reached it before convergence was reached (see the minimization.mdp file in the Zenodo deposit, https://zenodo.org/records/10973099).

Maximum forces of converged results were of the order of a couple hundreds $kJ.mol^{-1}.nm^{-1}$.

For CHARMM36m, systems were equilibrated for 2ns in the NVT ensemble with proteic atoms restrained with forces of 1000 $kJ.mol^{-1}.nm^{-1}$.

For the Amber forcefields, the equilibration was divided in two steps:

It turns out that the tleap program generates a water box with a density of around ~0.85.

This is usually resolved by the soft equilibration process recommended in the Amber protocol,[1] but creates a problem of pressure if one wants to directly perform the equilibration in NVT. We therefore performed a first step in the NPT ensemble for 200ps using the Berendsen thermostat so as to bring the density up to ~1.00 $kg.L^{-1}$.

We could then perform a 2ns equilibration in the NVT ensemble like for CHARMM36m.

Amber real-space cutoffs are not equivalent to CHARMM in terms of implementation.

CHARMM forcefields are calibrated with a switch of the forces so they smoothly decay to 0 $kJ.mol^{-1}.nm^{-1}$ as the cutoff is reached. On the other hand, Amber forcefields count on a shift of the potential and a correction of the energy. We therefore activated the vdw-modifier=Potential-shift-Verlet and the Dispcorr=EnerPres options. Cutoffs were then selected from the relevant papers for each one of the tested force-fields (Amber99SB-ILDN[2], AmberFF19SB[3], AmberFB18[4], CHARMM36m[5])

AmberFB18 was also tested on one replica for the entirely phosphorylated and non-phosphorylated cases with a real-space cutoff of 12 Å. An overall compaction of the peptide was observed, similar to what had been obtained using a 9 Å cutoff (see Figure 1 in the article).

### Dicussion on the simulations convergence

To assess the convergence of our simulations, we drew from Ellen Rieloff and Marie Skepö's work in the supplementary information of *Phosphorylation of a Disordered Peptide-Structural Effects and Force Field Inconsistencies,*[2] and used the radius of gyration (Rg) and end-to-end distance (Ree) as metrics to study convergence of our disordered peptides. Concepts and algorithms were implemented thanks to Alan Grossfield and Daniel M. Zuckermann's review *Quantifying uncertainty and sampling quality in biomolecular simulations*.[6] We report here only the general trends that can be extracted from convergence analysis of multiphosphorylated disordered peptides. All plots and scripts are available in the folder *Convergence* of the Zenodo repository: https://zenodo.org/records/12634115

#### Inter-replica convergence

Convergence is a difficult notion to define, especially for disordered systems. Although running several replicas is useful to explore different parts of the conformational landscape, finding common features between replicas can assess on the convergence and the quality of exploration of the trajectories. We therefore compare densities of Rg and Ree between the 3 replicas of each type of system.

1. R2 peptide

Generally speaking, all densities of replicas from the same type of system overlap rather substantially, which is a good sign of relative convergence between the replicas. Probably the most striking feature of the densities is that replicas of multiphosphorylated R2 seem to be much less alike than non-phosphorylated R2. This reveals the drastic modification of the conformational landscape induced by nP-collabs. A notable exception to this trend is indeed A19, which shows barely any difference between all types of replicas. The collapse of Rg and Ree for phosphorylated A18 and C36 below 12 angstroms is in complete disagreement with the wide profile of A19 and A99, attesting their opposing preference regarding nP-collabs.

1. Toy models

All distributions of Rg and Ree are extremely similar, with the exception of Replica 3 of peptide R+4 which displays a more collapsed state. Interestingly for all peptides but R+3, the Ree distributions reveal a low-populated state at 5 Å, corresponding to a conformation for which the terminal residues are directly interacting. All replicas seldom visit this state and are in qualitative agreement as to how often they do. We can therefore be confident that these trajectories have converged quite well.

**Intra-replica convergence**

A way to estimate whether convergence has been reached within a replica is to check whether the statistical error made on an observable has converged within the time of the simulation. We therefore implemented the block-averaging method as described in ref. (6) and applied it to each replica of our system. Briefly, the idea is to « cut » the trajectory in small blocks of size M, calculate the mean value of the observable in each block then calculate the standard error on this set of mean values : by repeating this calculation with increasing values of M, the standard error is supposed to asymptote towards the « real » standard error. We can therefore consider that a converged trajectory is a trajectory for which the standard error by block size asymptotes.

We ran the calculations from M=1 to M=5200 so that the latter block size still contains 10 elements per block.

1. R2 peptide

Asymptotic behavior seems to be reached for Rg in non-phosphorylated systems. Phosphorylated systems still exhibit a slightly increasing behavior of their error by block size. Surprisingly, A99 looks to be the less likely to have converged within 520 ns compared to the other forcefields, despite displaying very few nP-collabs.

The same observation between phosphorylated and non-phosphorylated systems can be made with Ree, with the notable exception of Replica 2 of C36 in sodium which does not converge as it adopts a beta-sheet conformation for half of the simulation (see DSSP profile).

2. Toy models

Convergence seems to be reached across all toy models for Rg, but not for all replicas of Ree. Error on Ree for Replica 3 of R+4 clearly continues to increase with block size, which is understandable since it is a replica where the 2P-collab ruptures for several dozens of nanoseconds. Interestingly, odd spacing numbers (R+1, R+3 and R+5) seem to converge towards a smaller error between all replicas compared to even spacing numbers (R+2, R+4 and R+6).

In conclusion, we can say that our simulations of R2 explore a vast amount of its conformational space but longer simulations will be required to fully sample the phosphorylated systems. This discrepancy between the non-phosphorylated and multiphosphorylated peptides seems to be related to the nP-collab phenomenon, which probably increases the stability of certain states, making the exploration more timely. The trajectories of the toy models look like they have fully converged, but the rupture of the 2P-collab in Replica 3 of R+4 indicates that there might be a dynamical property of nP-collabs at play with a timescale longer than the microsecond, at least with C36. We suggest that an interesting simulation to run would be a very long trajectory of the R+4 toy model to assess the lifetime of a nP-collab (we estimate from the breakage and reformation that at least 10$\mu$s would be required).

**Discussion on the ionic mobility**

The mobility of ions in the bulk and their interaction with water molecules were evaluated by running simulations of water boxes of 7 nm side-length with 31 $Cl^-$ ions and with 31 $K^+$ or $Na^+$ ions (corresponding to a concentration of 0.15mol.$L^{-1}$) , for each of the 4 chosen force-field/water combinations.

The Cumulative Distribution Function (CDF) is the integral of the pairwise Radial Distribution Fun ction (RDF), and can be interpreted as the average number of particles j at distance r from a particle i. We calculated the CDF of water oxygens with regard to cations in order to assess the coherency of our ion/water combinations of force-fields (Figure SI-112). Sodium and potassium behave in a remarkably similar fashion regardless of the force-field/water combination. This could be expected as the water coordination sphere is a common parameter for ionic parameterization. Of interest, we notice that the coordination sphere of $Na^+$ is more defined than the one of $K^+$ with a steeper plateau, which seems to indicate that $Na^+$ is more strongly coordinated to oxygens than $K^+$.

Frames were saved every 0.1ps and the self-part of the Van Hove function of each cation species was calculated with sampling rates of 0.1ps, 1ps, 10ps, 20ps, 50ps and 100ps (Figure SI-123).

The self-part of the Van Hove function corresponds to the probability of finding a particle i at time t+dt at a distance d from its position at time t. We noticed that for every force-field, potassium ions are more mobile than sodium ions in the fluid (higher mobility can be assessed by the self-part of the Van Hove shifting to higher distances). This could be explained by the more coordinated shell of sodium, which must hinder its capacity to diffuse freely.

The distributions become broader and centered around larger values when increasing the sampling rate. It is interesting to note that C36 ions are nearly twice as mobile as in the A18, A19 and A99 systems. This could have consequences on the formation of nP-collabs at low concentration of cations, since the probability to have a cation in position to be coordinated by phosphoresidues can be expected to increase with cation mobility in the solvent.

The most mobile ion type out of all combinations is the potassium of C36, which usually moves by around 4Å in 10ps (Figure SI-13a, right panel). We can therefore expect ions in all force-fields to move by less than the length of an amino-acid between two frames. Thus 10 ps sampling provides enough spatial resolution to capture the fine displacements of the ions in the fluid and in the vicinity of proteins, while remaining manageable in terms of data production. We therefore advise using a

10 ps sampling rate or lower for the study of ionic interactions with charged proteins if one is looking to discretely quantify ionic displacement.

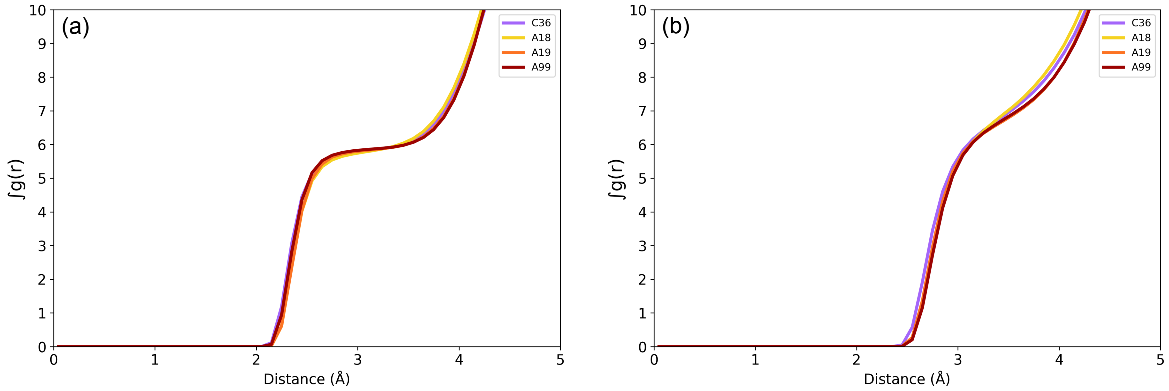


**Figure SI-12:** Cumulative distribution functions of water oxygens around cations as a function of the distance **(a)** $Na^+$ cations, **(b)** $K^+$ cations.

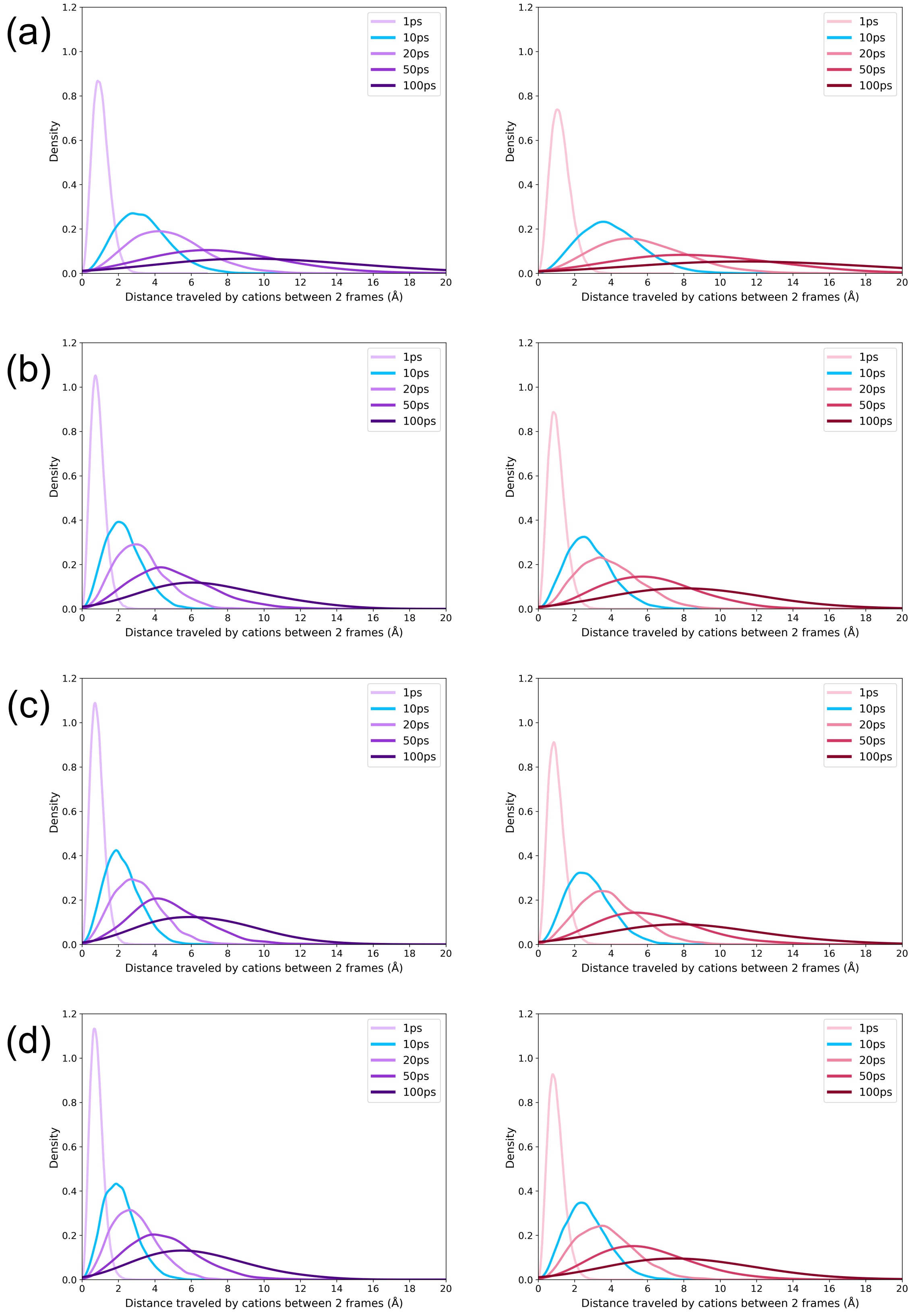


**Figure SI-13:** Cation mobility as a function of the trajectory sampling rate. Left panels: $Na^+$ cations, right panels: $K^+$ cations.

Force-field/water model combination : **(a)** C36, **(b)** A18, **(c)** A19, **(d)** A99
