## Supplementary material for "nP-collabs: Investigating counterion mediated bridges in the multiply phosphorylated tau-R2 repeat": AlphaFold3 structural prediction for the phosphorylated tau-R2 fragment: terms_of_use.md

# AlphaFold Server Output Terms of Use

BY USING ALPHAFOLD SERVER OUTPUT, YOU AGREE TO THE FOLLOWING TERMS OF USE.

By using structure predictions or related information provided by Google
DeepMind's [AlphaFold Server](https://alphafoldserver.com/) ("Output"), or
substantially derived from Output ("Derivatives"), without signing up to
AlphaFold Server and agreeing to
[AlphaFold Server Additional Terms of Service](https://alphafoldserver.com/terms),
you agree to be bound by these AlphaFold Server Output Terms of Use between you
(or your organization, as applicable) and Google LLC (these "Terms").

If you are using Output or Derivatives on behalf of an organization, you confirm
you are authorized either explicitly or implicitly to agree to, and are agreeing
to, these Terms as an employee or otherwise on behalf of your organization.

If you have a Google account and have agreed to
[AlphaFold Server Additional Terms of Service](https://alphafoldserver.com/terms),
your use of Output or Derivatives is governed by those terms. IF YOU HAVEN'T
AGREED TO
[ALPHAFOLD SERVER ADDITIONAL TERMS OF SERVICE](https://alphafoldserver.com/terms)
AND DON'T AGREE TO THESE TERMS, DON'T USE OUTPUT OR DERIVATIVES OR PERMIT ANY
THIRD PARTY TO DO SO ON YOUR BEHALF.

When we speak of "Google", "we" and "us", we mean Google LLC and its affiliates.

## Use restrictions

AlphaFold Server belongs to us. We make Output available free of charge, for
non-commercial use only, in accordance with following use restrictions. You must
not use nor allow others to use Output or Derivatives:

1.  ON BEHALF OF A COMMERCIAL ORGANIZATION OR IN CONNECTION WITH ANY
    COMMERCIAL ACTIVITIES, INCLUDING RESEARCH ON BEHALF OF A COMMERCIAL
    ORGANIZATION or sharing or using Output or Derivatives in a manner that
    will grant a commercial organization any rights in Output or Derivatives, in
    each case aside from indirectly via a scientific publication, open source
    release or to support journalism.
2.  IN CONNECTION WITH ANY AUTOMATED SYSTEM that predicts the binding or
    interaction of the protein with ligands or peptides, including, but not
    limited to, Glide or AutoDock.
3.  TO TRAIN MACHINE LEARNING MODELS OR RELATED TECHNOLOGY FOR BIOMOLECULAR
    STRUCTURE PREDICTION SIMILAR TO GOOGLE DEEPMIND'S
    [ALPHAFOLD](https://deepmind.google/technologies/alphafold/) TECHNOLOGY.
4.  WITHOUT PROVIDING CONSPICUOUS NOTICE THAT PUBLISHED OR DISTRIBUTED OUTPUT
    OR DERIVATIVES ARE PROVIDED UNDER AND SUBJECT TO THESE TERMS AND OF ANY
    MODIFICATIONS YOU MAKE TO OUTPUT.
    1.  If you remove these Terms, or any notice of these Terms, from Output or
        Derivatives, you must ensure further distribution is accompanied by a
        "Legally Binding Terms of Use" text file that contains the following
        notice: "This information is subject to AlphaFold Server Output Terms
        of Use found at
        [alphafoldserver.com/output-terms](https://alphafoldserver.com/output-terms)".
    2.  If you provide additional or different terms and conditions for use,
        reproduction or distribution of Output or Derivatives, you must include
        this "Use restrictions" section of these Terms as an enforceable
        provision and provide clear notice to subsequent users that Output and
        Derivatives are subject to such use restrictions.
5.  TO MISINFORM, MISREPRESENT OR MISLEAD, including:
    1.  misrepresenting your relationship with Google - including by using
        Google's trademarks, trade names, logos or suggesting endorsement by
        Google without Google's permission to do so - nothing in these Terms
        grants such permission;
    2.  misrepresenting the origin of Output or Derivatives;
    3.  distributing misleading claims of expertise or capability made
        particularly in sensitive areas (e.g., health); or
    4.  making decisions in domains that affect material or individual rights or
        well-being (e.g., healthcare).
6.  TO PERFORM OR FACILITATE DANGEROUS, ILLEGAL OR MALICIOUS ACTIVITIES,
    including promoting or facilitating the sale of, or providing instructions
    for synthesizing or accessing, illegal substances, goods or services.

Output and Derivatives may be provided with ancillary information that belongs
to us. You may use such ancillary information as allowed by these Terms. You
agree that your right to use and share Output or Derivatives is subject to your
compliance with these Terms. If you breach these Terms, Google reserves the
right to request that you delete and cease use or sharing of Output or
Derivatives in your possession or control. You agree to immediately comply with
any such request.

## Disclaimers and limitations of liability

Nothing in these Terms restricts any rights that cannot be restricted or limits
Google's responsibilities except as allowed by applicable law.

Structure predictions provided by AlphaFold Server are predictions with varying
levels of confidence, provided strictly as-is and without representation or
warranty of any kind, express or implied, including implied warranties of
merchantability, fitness for a particular purpose, title, and non-infringement.
You are responsible for the accuracy, reliability, availability, effectiveness,
or correct use of information generated by AlphaFold Server or for any impact
from ongoing use of such information. If you rely on any such information, you
do so solely at your own risk.

AlphaFold Server, Output and Derivatives are for theoretical modeling only.
These are not intended, validated, or approved for clinical use. You should not
use these for clinical purposes or rely on them for medical or other
professional advice. Any content regarding those topics is provided for
informational purposes only and is not a substitute for advice from a qualified
professional.

In no circumstances will Google be responsible for any indirect, special,
incidental, exemplary, consequential, or punitive damages, or lost profits of
any kind or any liability, including for its own negligence, relating to these
Terms over $500.

## Governing law and disputes

These Terms will be governed by the laws of the State of California without
regard to choice of law principles. The state or federal courts of Santa Clara
County, California shall have exclusive jurisdiction of any dispute arising out
of these Terms.

Given the nature of scientific research, it may take some time for any breach of
these Terms to become apparent. To the extent allowed by applicable law, any
legal claims relating to these Terms can be initiated until the later of: (a)
the cut-off date under applicable law for bringing the legal claim; or (b) two
years from the date you or Google (as applicable) became aware, or should
reasonably have become aware, of the facts giving rise to that claim. You won't
argue limitation, time bar, delay, waiver or the like in an attempt to bar an
action filed within that time period, and neither will we.

## Miscellaneous

Google may update these Terms (1) to reflect changes in how it does business,
(2) for legal, regulatory, or security reasons, or (3) to prevent abuse or harm.
The version of these Terms that were effective on the date the relevant Output
was generated will apply to your use of that Output or its Derivatives.

If it turns out that a particular provision of these Terms is not valid or
enforceable, this will not affect any other provisions.
